# Cryo-EM visualizes multiple steps of dynein’s activation pathway

**DOI:** 10.1101/2024.09.28.615567

**Authors:** Agnieszka A. Kendrick, Kendrick H. V. Nguyen, Wen Ma, Eva P. Karasmanis, Rommie E. Amaro, Samara L. Reck-Peterson, Andres E. Leschziner

## Abstract

Cytoplasmic dynein-1 (dynein) is an essential molecular motor controlled in part by autoinhibition. We recently identified a structure of partially autoinhibited dynein bound to Lis1, a key dynein regulator mutated in the neurodevelopmental disease lissencephaly. This structure provides an intermediate state in dynein’s activation pathway; however, other structural information is needed to fully explain Lis1 function in dynein activation. Here, we used cryo-EM and samples incubated with ATP for different times to reveal novel conformations that we propose represent intermediate states in the dynein’s activation pathway. We solved sixteen high-resolution structures, including seven distinct dynein and dynein-Lis1 structures from the same sample. Our data also support a model in which Lis1 relieves dynein autoinhibition by increasing its basal ATP hydrolysis rate and promoting conformations compatible with complex assembly and motility. Together, this analysis advances our understanding of dynein activation and the contribution of Lis1 to this process.

---

Cytoplasmic dynein-1 (dynein) is a highly conserved minus-end directed microtubule motor that transports numerous cargoes, including RNAs, membrane vesicles, and viruses, and plays a critical role in cell division^1^. The dynein complex (1.4 MDa) comprises a dimer of two motor domains and two copies each of five accessory chains (an intermediate chain, a light intermediate chain, and three light chains)^2^. Cellular and structural work suggests that dynein exists largely in an autoinhibited ‘Phi’ conformation in cells^3–5^. Upon relief from autoinhibition, dynein undergoes conformational changes to assemble into an active complex (∼ 4 MDa) with its cofactors: a multisubunit dynactin complex (1.1 MDa) and a coiled-coil activating adaptor that also links the active complex to cargo^1,6^. Mutations in dynein and its binding partners are linked to neurodevelopmental and neurodegenerative diseases^1,7^.

Dynein is a member of the AAA+ (<u>A</u>TPase <u>a</u>ssociated with various cellular <u>a</u>ctivities) family of proteins. Its “heavy chain” includes the motor domain composed of six AAA+ modules, a long stalk, a microtubule-binding domain (MTBD) located at the end of the stalk, a “buttress” region that makes additional connections between the AAA5 module and the stalk, and a long tail that interacts with dynactin and an activating adaptor (Fig. 1a). The tail is also responsible for the generation of a power stroke through its mechanical element, called the “linker”^8^. Out of the six AAA+ modules, four bind ATP (AAA1, AAA2, AAA3, and AAA4) but only three hydrolyze it (AAA1, AAA3, and AAA4)^9–13^. AAA5 and AAA6 do not contain the residues necessary for nucleotide binding^11,13^. Each AAA module is composed of a large and small domain and nucleotide binding takes place in a groove between them and the large domain of the adjacent AAA module (Fig. 1b).

**Figure 1.**
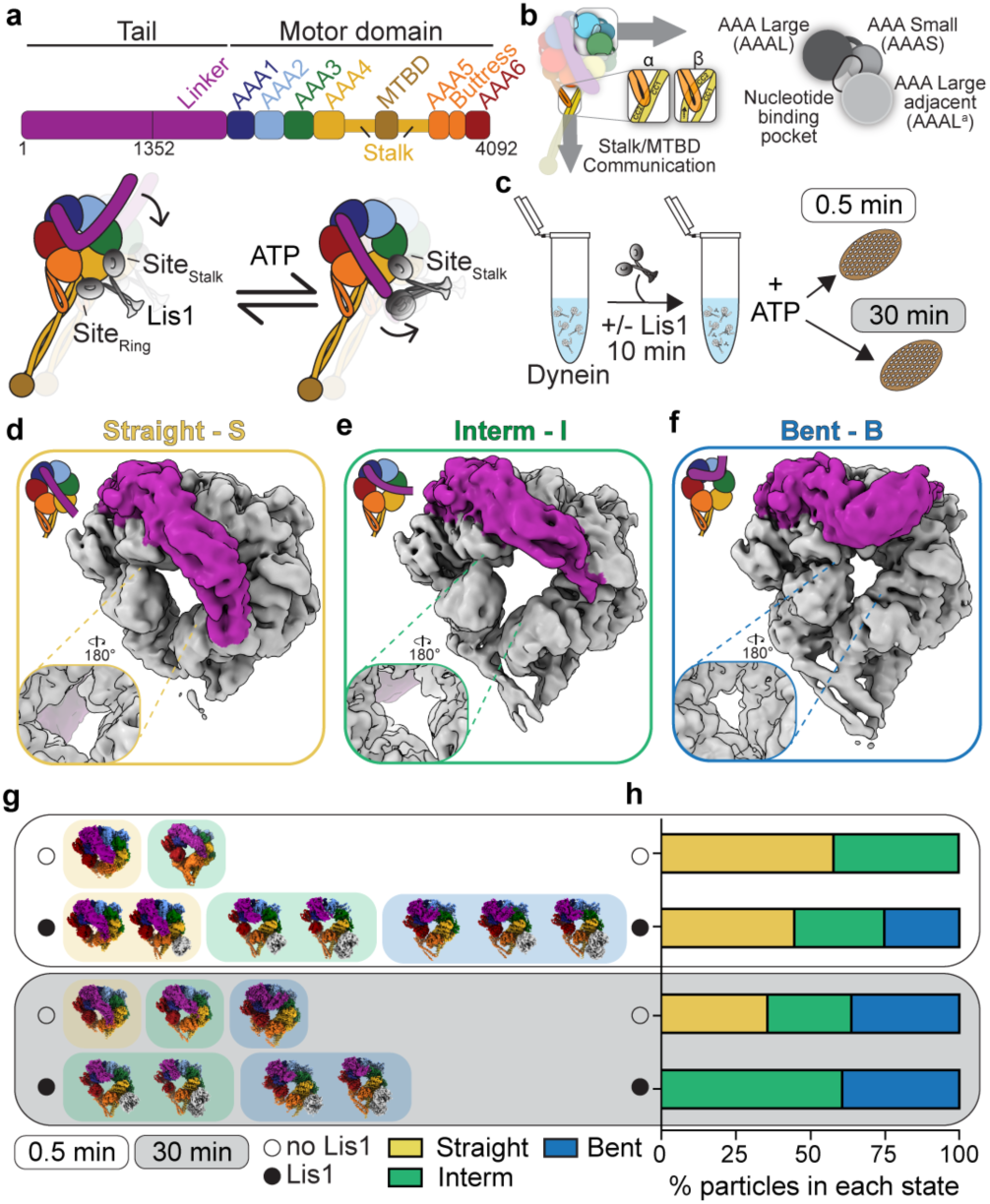
Time-resolved cryo-EM captures Lis1’s effect on dynein’s conformational landscape during ATP hydrolysis. **a.** Cartoon of dynein domain organization. AAA – <u>A</u>TPase <u>a</u>ssociated with various cellular <u>a</u>ctivities, MTBD – microtubule-binding domain. Lis1 binding at different sites on dynein (sitestalk and sitering) is shown in different dynein conformations^41^. **b.** The architecture of a nucleotide binding pocket and Stalk/MTBD communication. **c.** Experimental setup. **d. – f.** The three major groups of conformations identified in the datasets are shown on locally refined cryo-EM volumes filtered to 6 Å: **d.** straight linker (S) – yellow outline, **e.** intermediate (Interm, I) linker – green outline, and **f.** bent linker (B) – blue outline. The linker region (amino acids 1-1772) is highlighted in magenta. The extent of ring opening is shown to the left of each panel. **g.** Different conformations were identified in each dataset in the absence (open circle) or presence of Lis1 (black circle) at two different time points (0.5 min – white background and 30 min – gray background). **h.** Relative abundance of particles belonging to different states obtained from particle distributions in cryo-EM datasets in the absence (open circle) or presence of Lis1 (black circle) at two different time points (0.5 min – white background and 30 min – gray background).

Dynein converts the chemical energy of ATP into movement and force through its mechanochemical cycle^14^. During it, ATP binding and hydrolysis in the primary catalytic module, AAA1, is linked to large conformational changes that lead to the opening and closing of the AAA ring, rearrangement of the linker to generate the power stroke, and reorganization of helices in the buttress and stalk (CC1 and CC2) to regulate dynein’s interaction with microtubules via the MTBD (Fig. 1b and c)^10,15,16^. This reorganization is due to the sliding of the CC1 with respect to CC2. The two main “registers” that the helices can adopt, referred to as ⍺ and β, correspond to high and low microtubule binding affinities, respectively (Fig. 1b)^17,18^. Mutations in AAA3 that prevent it from hydrolyzing ATP also slow down ATP hydrolysis in AAA1, providing evidence for an allosteric role for AAA3 in regulating dynein’s mechanochemistry^10,16,19^.

Dynein function in vivo is tightly linked to Lis1, the only dynein regulator that binds directly to dynein’s motor domain^20,21^. Lis1 is a dimer initially identified as a gene (*PAFAH1B1*) mutated in the neurodevelopmental disease lissencephaly, also linked to mutations in dynein’s heavy chain (*DYNC1H1*)^22–24^. Lis1 is conserved from fungi, including *S. cerevisiae*, to humans. Dynein is important for mitotic spindle positioning in yeast, though it is non-essential in that organism, providing an excellent model system to study dynein function^25–29^. Like human dynein, yeast dynein requires dynactin and the presumed activating adaptor Num1 for function^26,27,30^. Work in yeast and other organisms has shown that Lis1 is important for forming active dynein complexes^31–35^. Recently, we showed that Lis1 does so by relieving dynein’s autoinhibited “Phi” conformation. We found that two Lis1 dimers could disrupt the Phi conformation by wedging themselves between two dynein protomers (an intermediate state called “Chi”), potentially priming dynein for subsequent complex assembly and activation^36^. Lis1 is also involved in further steps of the dynein activation pathway by directly interacting with dynactin to support conformations that allow complex assembly^37^. In addition, dynein-dynactin-activating adaptor complexes assembled in the presence of Lis1 recruit an additional dynein dimer, and these complexes move faster on microtubules^31,33^. Taken together, these findings suggest multiple roles for Lis1 in regulating dynein function that may go beyond Phi particle opening. Indeed, mutations in dynein that block the formation of Phi do not fully rescue Lis1 disruption phenotypes^35,36,38,39^. Thus, more structural and mechanistic studies are needed to understand how Lis1 regulates dynein.

Lis1 binds to dynein at three sites, one between AAA3 and AAA4 (site_ring_), one in the stalk region (site_stalk_) (Fig. 1a), and a third site that is accessible in the presence of a second dynein protomer in the Chi conformation, in trans at AAA5/ AAA6^21,32,40,41^. Previous analysis showed that the nucleotide state at AAA3 regulates Lis1’s association with dynein and dynein’s microtubule-binding state^40^. However, recently identified roles of Lis1 in dynein activation and complex assembly point to a mechanism by which Lis1 regulates dynein prior to dynein’s interaction with microtubules. Indeed, a dynein mutant locked in a microtubule-unbound state binds to Lis1 with a stronger affinity than its microtubule-bound counterpart, consistent with data showing that Lis1 can dissociate from active dynein complexes bound to microtubules^31,33,42^. In addition, a cryo-EM structure of two dynein dimers complexed with dynactin and an activating adaptor (JIP3) revealed that Lis1 bound to the dynein dimer that was detached from microtubules, while the microtubule-bound dynein dimer was not associated with Lis1^37^. Although the authors also suggested that Lis1 changes the nucleotide state of the AAA3 domain, it is unclear what role Lis1 plays in the regulation of dynein’s motor activity. Numerous lines of evidence suggest that Lis1 does not affect dynein’s microtubule-stimulated ATPase activity, but how Lis1 affects dynein’s basal ATPase activity, i.e. when dynein is not bound to microtubules, and during dynein activation remains unclear^20,21,41,43,44^.

To address these knowledge gaps in our understanding of dynein’s mechanochemistry and its regulation by Lis1, we used cryo-EM combined with a time-resolved sample preparation pipeline in the presence of ATP to capture as many states as we could detect of dynein throughout its mechanochemical cycle (Fig. 1c). Using wild-type dynein monomer and wild-type Lis1 from *S. cerevisiae* we solved 16 unique dynein and dynein/ Lis1 structures, including previously unknown states. Our data show that Lis1 is affecting dynein’s conformational landscape by increasing its basal ATPase rate, a finding previously reported but not fully explored^21,41^. Our data support a hypothesis in which Lis1 promotes dynein conformations that allow for a transition from an autoinhibited state to an open conformation by facilitating a nucleotide exchange in AAA3, potentially priming dynein for complex assembly. As this model predicts, opening the Phi particle via Lis1 binding or by mutating a dynein residue important for Phi formation (D2868K) increases dynein’s basal ATPase rate^38^. Taken together, our analysis suggests that dynein’s ATPase activity is important for its activation, and that, by binding to dynein, Lis1 assists dynein’s transition into a state primed for subsequent complex assembly with dynactin and an activating adaptor.

## Cryo-EM shows different conformations during dynein’s mechanochemical cycle and Lis1 binding

Previous high-resolution structures of dynein in complex with Lis1 described the interactions between these proteins. However, these structures were solved using mutations or non-hydrolyzable ATP analogs that limited the ability to capture dynein’s structural dynamics^32,36,40,45^. To characterize the conformational landscape of dynein during Lis1 binding and ATP hydrolysis, we employed a time-resolved freezing approach. We purified wild-type yeast dynein monomer (amino acids 1352-4092) in the absence of any nucleotide and plunge froze grids after the addition of excess ATP (500-fold) at two different time points (0.5 min and 30 min), incubating the samples at 4°C to slow down nucleotide hydrolysis. We also prepared grids with samples that were preincubated with Lis1 prior to ATP addition (Fig. 1c). After data collection and analysis (Extended Data Fig. 1 – 4, and Table 1 and 2), we observed the expected high level of heterogeneity in our datasets. We grouped the reconstructions into three distinct conformations based on the position of dynein’s linker, a mechanical element that adopts different conformations in response to the ATP hydrolysis-driven opening and closing of the ring: “straight” – S, yellow outline (Fig. 1d), “interm” (intermediate) – I, green outline (Fig. 1e), and “bent” – B, blue outline (Fig. 1f). We named all our maps with letters corresponding to the different conformational states and numbers starting with the shorter time point (0.5 min) (Extended Data Fig. 1 – 4 and Table 1 and 2).

**Table 1:** Cryo-EM data information and model validation statistics for the 0.5 min datasets. CryoEM data collection parameters, reconstruction information, and model refinement statistics.

| Name | S1 | S2 | S3 | I1 | I2 | I3 | B1 | B2 | B3 | B4 |
| --- | --- | --- | --- | --- | --- | --- | --- | --- | --- | --- |
| State | Straight | Straight | Straight /Lis1 | Interm. /Lis1 | Interm. /1xLis1 | Interm. /2xLis1 | Bent /Lis1 | Bent | Bent /1xLis1 | Bent /2xLis1 |
| Condition (Lis1) | n/a | Lis1 | Lis1 | Lis1 | Lis1 | Lis1 | Lis1 | Lis1 | Lis1 | Lis1 |
| Time point (min) | 0.5 | 0.5 | 0.5 | 0.5 | 0.5 | 0.5 | 0.5 | 0.5 | 0.5 | 0.5 |
| EMD- | 46919 | 46897 | 46938 | 47019 | 46958 | 46941 | 46935 | 46940 | 46942 | 46919 |
| PDB- | 9DIU | 9DI3 | 9DJU | 9DMW | 9DKH | 9DJZ | 9DKM | 9DJY | 9DK0 | 9DJ7 |
| Data collection and processing |  |  |  |  |  |  |  |  |  |  |
| Magnification | 36 000 |  |  |  |  |  |  |  |  |  |
| Voltage (kV) | 200 |  |  |  |  |  |  |  |  |  |
| Electron exposure (e−/Å²) | Dataset #1 – 54<br>Dataset #2 – 54<br>Dataset #3 – 55 | Dataset #1 – 53<br>Dataset #2 – 54<br>Dataset #3 – 54<br>Dataset #4 – 51 |  |  |  |  |  |  |  |  |
| Defocus range (μm) | 0.8 – 2.4 |  |  |  |  |  |  |  |  |  |
| Pixel size (Å) | 1.16 |  |  |  |  |  |  |  |  |  |
| Symmetry imposed | C1 |  |  |  |  |  |  |  |  |  |
| Images (no.) | 1 849 | 12 370 |  |  |  |  |  |  |  |  |
| Initial particles (no.) | 319 552 | 1 047 074 |  |  |  |  |  |  |  |  |
| Final particles (no.) | 46 585 | 68 450 | 68 486 | 105 604 | 46 266 | 33 037 | 92 164 | 23 426 | 40 468 | 25 170 |
| Map resolution (Å) (FSC 0.143) | 4 | 4.1 | 4 | 3.7 | 3.9 | 3.9 | 3.3 | 3.6 | 3.7 | 3.8 |
| Refinement |  |  |  |  |  |  |  |  |  |  |
| Initial model used (PDB code) | 7MGM | 7MGM | 7MGM | 7MGM | 7MGM | 7MGM | 7MGM | 7MGM | 7MGM | 7MGM |
| Model resolution (Å) (FSC 0.5) | 4.5 | 4.3 | 4.4 | 4 | 4.2 | 4.2 | 3.4 | 3.9 | 3.9 | 4 |
| Map sharpening *B* factor (Å²) | 86.5 | 106 | 109.8 | 97.4 | 85.1 | 67.4 | 83.9 | 63 | 72.6 | 60.8 |
| Model composition |  |  |  |  |  |  |  |  |  |  |
| Non-hydrogen atoms | 17 847 | 17 694 | 20 676 | 21 212 | 21 004 | 23 913 | 21 339 | 18 424 | 21 050 | 23 851 |
| Protein residues | 2302 | 2 282 | 2 643 | 2 729 | 2691 | 3047 | 2 748 | 2 341 | 2 690 | 3026 |
| Ligands | ADP:3, ATP:1 | ADP:3, ATP:1 | ADP:3, ATP:1 | MG:1, ADP:3, ATP:1 | MG:1, ADP:3, ATP:1 | MG:1, ADP:3, ATP:1 | MG:1, ADP:3, ATP:1 | MG:1, ADP:3, ATP:1 | MG:1, ADP:3, ATP:1 | MG:1, ADP:3, ATP:1 |
| *B* factors (Å²) |  |  |  |  |  |  |  |  |  |  |
| Protein | 166.53 | 124.09 | 140.15 | 69.90 | 107.18 | 141.67 | 54.14 | 98.06 | 71.21 | 94.99 |
| Ligand | 162.06 | 92.88 | 70.03 | 43.58 | 69.38 | 118.25 | 25.38 | 80.48 | 46.11 | 76.40 |
| R.m.s. deviations |  |  |  |  |  |  |  |  |  |  |

|  |  |  |  |  |  |  |  |  |  |  |
| --- | --- | --- | --- | --- | --- | --- | --- | --- | --- | --- |
| Bond lengths (Å) | 0.003 | 0.004 | 0.003 | 0.003 | 0.003 | 0.003 | 0.003 | 0.003 | 0.003 | 0.003 |
| Bond angles (°) | 0.715 | 0.680 | 0.740 | 0.701 | 0.647 | 0.616 | 0.562 | 0.585 | 0.602 | 0.613 |
| Validation |  |  |  |  |  |  |  |  |  |  |
| MolProbity score | 1.51 | 1.75 | 1.68 | 1.40 | 1.52 | 1.55 | 1.37 | 1.30 | 1.44 | 1.51 |
| Clashscore | 7.75 | 9.34 | 8.46 | 5.16 | 6.31 | 7.23 | 4.93 | 4.81 | 5.96 | 6.48 |
| Poor rotamers (%) | 0.3 | 0.2 | 0.5 | 0.1 | 0 | 0.1 | 0.1 | 0.1 | 0.1 | 0.2 |
| Ramachandran plot |  |  |  |  |  |  |  |  |  |  |
| Favored (%) | 97.60 | 96.19 | 96.55 | 97.35 | 97.04 | 97.18 | 97.45 | 97.75 | 97.41 | 97.15 |
| Allowed (%) | 2.4 | 3.81 | 3.45 | 2.65 | 2.96 | 2.82 | 2.55 | 2.25 | 2.59 | 2.85 |
| Disallowed (%) | 0 | 0 | 0 | 0 | 0 | 0 | 0 | 0 | 0 | 0 |

**Table 2:** Cryo-EM data information and model validation statistics for the 30 min datasets. CryoEM data collection parameters, reconstruction information, and model refinement statistics.

| Name | S4 | I4 | B5 | I5 | I6 | I7 | B6 | B7 | B8 |
| --- | --- | --- | --- | --- | --- | --- | --- | --- | --- |
| State | Straight | Interm | Bent | Interm /Lis1 | Interm /1xLis1 | Interm /2xLis1 | Bent /Lis1 | Bent /1xLis1 | Bent /2xLis1 |
| Condition (Lis1) | n/a | n/a | n/a | Lis1 | Lis1 | Lis1 | Lis1 | Lis1 | Lis1 |
| Time point (min) | 30 | 30 | 30 | 30 | 30 | 30 | 30 | 30 | 30 |
| EMD- | 46953 | 46954 | 46959 | 46974 | 46972 | 46975 | 47033 | 47026 | 47032 |
| PDB- | 9DKD | 9DKE | 9DKJ | 9DLD | 9DKX | 9DLE | 9DNB | 9DN5 | 9DN7 |
| Data collection and processing |  |  |  |  |  |  |  |  |  |
| Magnification | 150 000 |  |  |  |  |  |  |  |  |
| Voltage (kV) | 200 |  |  |  |  |  |  |  |  |
| Electron exposure (e−/Å²) | 54 |  |  | 54 |  |  |  |  |  |
| Defocus range (μm) | 0.75 – 2.8 |  |  |  |  |  |  |  |  |
| Pixel size (Å) | 0.889 |  |  |  |  |  |  |  |  |
| Symmetry imposed | C1 |  |  |  |  |  |  |  |  |
| Images (no.) | 7 227 |  |  | 6 670 |  |  |  |  |  |
| Initial particles (no.) | 1 047 074 |  |  | 744 012 |  |  |  |  |  |
| Final particles (no.) | 119 768 | 94 448 | 122 388 | 106 139 | 59 650 | 40 384 | 67 155 | 35 055 | 28 077 |
| Map resolution (Å) (FSC 0.143) | 3.3 | 3.6 | 2.8 | 3.2 | 3.4 | 3.4 | 3 | 3.3 | 3.25 |
| Refinement |  |  |  |  |  |  |  |  |  |
| Initial model used (PDB code) | 7MGM | 7MGM | 7MGM | 7MGM | 7MGM | 7MGM | 7MGM | 7MGM | 7MGM |
| Model resolution (Å) (FSC 0.5) | 3.6 | 4.2 | 3.0 | 3.4 | 3.7 | 3.7 | 3.3 | 3.7 | 3.5 |
| Map sharpening *B* factor (Å²) | 82.3 | 74.3 | 71.2 | 77.2 | 70.1 | 56.5 | 57.1 | 55.1 | 51.8 |
| Model composition |  |  |  |  |  |  |  |  |  |
| Non-hydrogen atoms | 17 901 | 17 457 | 18 075 | 22 996 | 20 808 | 22 893 | 22 963 | 21000 | 23772 |
| Protein residues | 2 313 | 2 291 | 2 291 | 3 017 | 2646 | 2985 | 3020 | 2679 | 3021 |
| Ligands | MG:1, ADP:3, ATP:1 | MG:1, ADP:3, ATP:1 | MG:2, ADP:3, ATP:1 | MG:3, ADP:3, ATP:1 | MG:3, ADP:3, ATP:1 | MG:2, ADP:3, ATP:1 | MG:1, ADP:3, ATP:1 | MG:1, ADP:3, ATP:1 | MG:1, ADP:3, ATP:1 |
| *B* factors (Å²) |  |  |  |  |  |  |  |  |  |
| Protein | 67.01 | 123.03 | 66.14 | 96.14 | 106.31 | 108.74 | 69.30 | 99.61 | 84.55 |
| Ligand | 30.25 | 86.62 | 39.03 | 55.26 | 69.26 | 71.52 | 32.12 | 69.54 | 62.99 |
| R.m.s. deviations |  |  |  |  |  |  |  |  |  |
| Bond lengths (Å) | 0.003 | 0.003 | 0.003 | 0.003 | 0.003 | 0.003 | 0.002 | 0.003 | 0.003 |

|  |  |  |  |  |  |  |  |  |  |
| --- | --- | --- | --- | --- | --- | --- | --- | --- | --- |
| Bond angles (°) | 0.655 | 0.715 | 0.628 | 0.597 | 0.621 | 0.600 | 0.521 | 0.623 | 0.579 |
| **Validation** |  |  |  |  |  |  |  |  |  |
| MolProbity score | 1.50 | 1.43 | 1.19 | 1.48 | 1.46 | 1.49 | 1.37 | 1.35 | 1.52 |
| Clashscore | 7.00 | 6.2 | 4.06 | 6.19 | 5.41 | 5.76 | 6.71 | 6.43 | 6.17 |
| Poor rotamers (%) | 0.1 | 0.1 | 0.1 | 0.1 | 0.1 | 0.2 | 0 | 0 | 0 |
| **Ramachandran plot** |  |  |  |  |  |  |  |  |  |
| Favored (%) | 97.44 | 97.58 | 98.01 | 97.27 | 97.05 | 97.01 | 98.25 | 98.49 | 96.91 |
| Allowed (%) | 2.56 | 2.42 | 1.99 | 2.73 | 2.95 | 2.99 | 1.75 | 1.51 | 3.09 |
| Disallowed (%) | 0 | 0 | 0 | 0 | 0 | 0 | 0 | 0 | 0 |

To understand how Lis1 binding affects dynein’s conformational landscape, we next examined the number of total particles that belong to each conformation (bent – blue, intermediate – green, and straight – yellow) in each dataset collected at the two different timepoints and in the presence or absence of Lis1 (Fig. 1g). This analysis showed that dynein alone samples more states when the incubation is extended from 0.5 min to 30 min (Fig. 1g and h). However, the addition of Lis1 increases the number of states dynein samples in the shorter time point (0.5 min) (Fig. 1g and h). To exclude the possibility that these differences could reflect the different sizes of datasets, we repeated the analysis by combining data belonging to the same experimental condition (+Lis1 or -Lis1) but were collected at different time points (Extended Data Fig. 5a and b). Because these datasets were collected using different microscopes, we only processed them together until we obtained initial 3D reconstructed models, without any prior structural information. We then mapped the particles of each model to their corresponding datasets and conformational subgroups (bent, intermediate, and straight linker). This analysis showed a similar increase in the number of conformational states seen in the presence of Lis1 at a shorter time point as the one we observed when each dataset was processed separately (Extended Data Fig. 5c). This suggests that Lis1 may be facilitating dynein’s transition between different conformations, allowing dynein to sample more states. Next, we will discuss the different dynein conformations we observed in the absence and presence of Lis1, followed by how these conformations might explain the effects of Lis1 on dynein’s mechanochemistry.

## Dynein’s conformational landscape during ATP hydrolysis

To understand how ATP hydrolysis affects dynein structurally, we built models for the different high-resolution dynein conformations identified in our datasets and mapped nucleotides to their binding pockets, as all our structures showed nucleotide densities in the AAA modules (Fig. 2a, b, and c, Extended Data Fig. 4a – S1 and S4, 4b – I4, 4c – B2, and, and Extended Data Fig. 6a). In each linker state (bent, intermediate, and straight) dynein is bound to ADP in AAA1 and AAA3, although in some states there is also density for Mg^2+^, indicating a tighter binding state, as Mg^2+^ release weakens ADP binding^46,47^. Although our maps only have partial densities in the stalk region, the ring opening and the stalk’s CC1 and CC2 positions in dynein with a straight linker is consistent with a strong microtubule binding state, as indicated by the lack of bulging of CC2 in the stalk region characteristic of the weak microtubule binding state (Fig. 2a – Stalk vs. 2c – Stalk)^48^. The intermediate linker dynein has partial density that fits a model in which the CC2 has a slight bulge in the stalk and is consistent with a semi-⍺ or semi-bent state that could represent an intermediate microtubule binding state (Fig. 2b – Stalk)^18,49,50^. In agreement with having occupied nucleotide binding pockets, the ring of the straight linker dynein is more closed than in the crystal structure of apo-dynein (PDB: 4AI6, Fig. 2d). We see a partial closing of the AAA1 binding pocket, indicative of a post-hydrolysis ADP-bound sate (Fig. 2e), and ADP and Mg^2+^ in AAA3 in both straight and intermediate dynein as opposed to ADP without Mg^2+^ in the bent linker dynein (Fig. 2a) or crystal structure (PDB: 4AI6, Fig. 2f)^15^.

**Figure 2.**
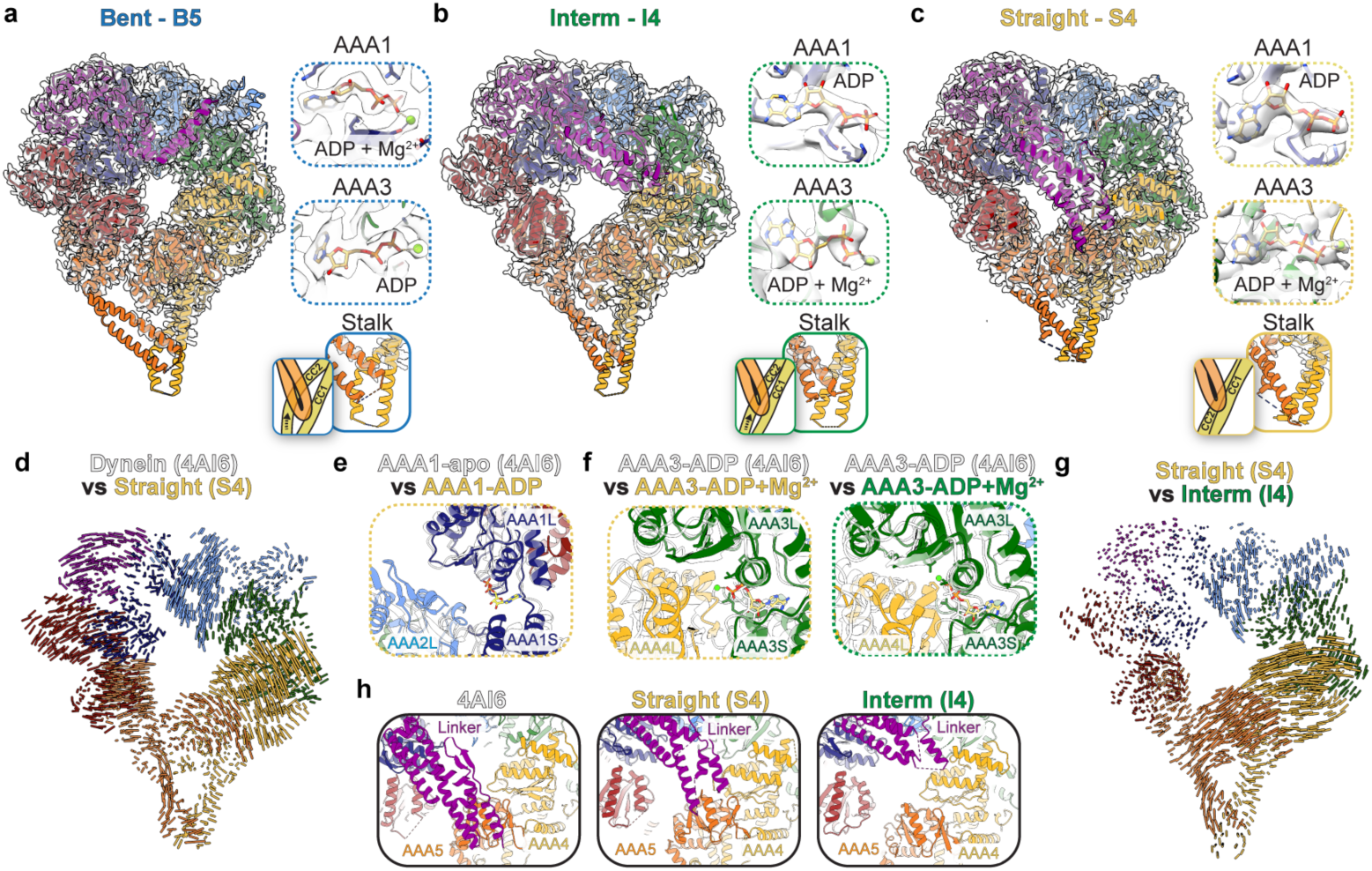
Conformational landscape of dynein during ATP hydrolysis. **a.** Models of dynein for the bent – B5 (**a**), intermediate – I4 (**b**), and straight – S4 (**c**) conformations from the 30 min with ATP dataset fit into their corresponding cryo-EM density maps. Each structural element is colored based on the dynein schematic in Fig. 1a. To the right of each fitted model are close ups of the nucleotide binding pockets for AAA1 and AAA3, and of the predicted stalk conformations. **d**. Map of pairwise alpha-carbon distances between straight linker model (S4) and an X-ray structure of *S. cerevisiae* dynein in AAA1-apo (PDB: 4AI6), with the models aligned relative to their AAA1 modules. The length of each vector is proportional to the interatomic distance. Residues 1361-1772 of the linker were removed for clarity. **e.** Comparison of AAA1 nucleotide binding pockets for straight linker dynein (S4) bound to ADP with *S. cerevisiae* in AAA1-apo (PDB: 4AI6). The models were aligned based on residues 1797-1894 in AAA1. **f.** Comparison of AAA3 nucleotide binding pockets for straight linker dynein (S3, left panel – yellow outline) and intermediate linker dynein (I4, right panel – green outline) with *S. cerevisiae* (PDB: 4AI6). The models were aligned based on residues 2377-2445 in AAA3. **g.** View of linker docking in PDB: 4AI6 (left panel), straight linker conformation (S4, middle panel), and intermediate linker conformation (I4, right panel). **h.** Map of pairwise alpha-carbon distances between the straight linker (S4) and the intermediate linker (I4) models. The length of each vector is proportional to the interatomic distance. The models were aligned relative to their AAA1 modules. Residues 1361-1772 of the linker were removed for clarity.

The intermediate linker dynein states identified in our datasets have similar conformations to the ones identified in a recent analysis of full length human dynein^51^. In our dataset, the comparison of the straight and intermediate linker dynein shows movement in the AAA domains, even though the nucleotide states are the same in these structures (Fig. 2g, and 2b and 2c). While the linker docks at AAA5 in the apo (PDB: 4AI6) and straight linker dynein, the linker docks at AAA4 in the intermediate linker dynein (Fig. 2h). These differences in the ring and linker without corresponding changes in the nucleotide state suggest some flexibility in dynein that is independent of ATP hydrolysis and is consistent with previous FRET analysis and recent human dynein structures^11,51^.

## The effect of Lis1 on dynein’s conformational landscape

To understand how Lis1 affects dynein’s mechano- chemistry, we next analyzed the datasets where dynein had been preincubated with Lis1 before the addition of ATP. These datasets also showed high level of heterogeneity (Supplemental Video 1) with the bent and intermediate linker dynein states in the 0.5 min and 30 min conditions (Fig. 3a and 3b, and Extended Data Fig. 6b and 6c) while the straight linker states where only detected in the 0.5 min condition (Fig. 1g and 1h, 3c and 3d, and Extended Data Fig. 6b). In addition, the nucleotide occupancies and stalk conformations in these different dynein states are similar to those seen in the absence of Lis1 (Fig. 2a – c and 3a – d, Extended Data Fig. 4 and 6).

**Figure 3:**
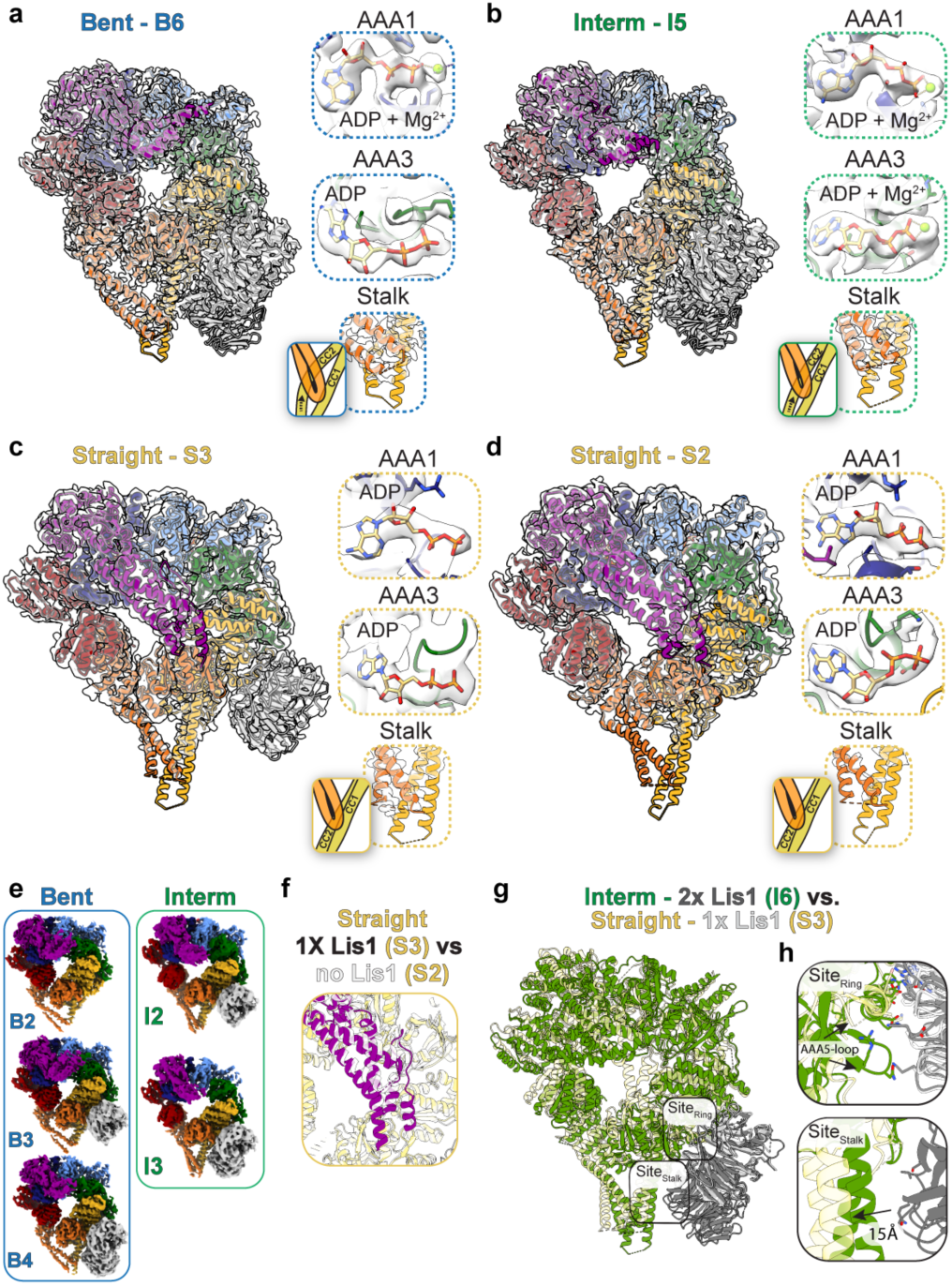
Lis1 binding to dynein expands dynein’s conformational landscape. Models of dynein bound to Lis1 for: the bent – B6 (**a**) and intermediate – I5 (**b**) dynein from the same dataset (dynein with Lis1 incubated for 30 min with ATP), and straight with Lis1 bound – S3 (**c**) and straight with no Lis1 bound – S2 (**d**) dynein from the same dataset (dynein with Lis1 incubated for 0.5 min with ATP), fit into their corresponding cryo-EM density maps. Each structural element is colored based on the dynein schematic in Fig. 1a. To the right of each fitted model are views of the nucleotide binding pockets for AAA1 and AAA3 and of the stalk conformations. **e**. Additional cryo-EM maps obtained after heterogeneity analysis of bent (blue box) and intermediate (green box) dynein conformations from the dynein with Lis1 incubated for 0.5 min with ATP dataset after 3D classification show one or two Lis1 bound to dynein. **f**. Comparison of linker (magenta) in straight linker dynein (S3, yellow) bound to Lis1 with linker (white) in straight linker dynein (S2, white) with no Lis1 bound. The models were aligned based on the position of AAA1. **g**. Comparison of models built for intermediate state dynein (I6, green) bound to 2 Lis1 β-propellers (gray) with straight linker dynein (S3, yellow) bound to one Lis1 β-propeller (white). The two Lis1 binding sites are highlighted. The models were aligned based on the position of Lis1 bound to sitering. **h**. The dynein/ Lis1 binding sites from g.

Because some of our maps showed partial density for the Lis1 β-propellers, we performed 3D classifications without alignment in Relion to further resolve this compositional heterogeneity (Extended Data Fig. 2 and 3). This analysis separated states where we see no, one, or two Lis1 β-propellers bound to dynein (Fig. 3e and Extended Data Fig. 2 – 4, and 6d and e). The nucleotide occupancies in these new maps correspond to the nucleotides observed in pre-3D classification maps (Fig. 3a – d, Extended Data Fig. 4 and 6). In addition, the interaction sites between Lis1 and dynein in our maps are those previously characterized and are largely the same in the bent and intermediate dynein states (Extended Data Fig. 7a – c).

## Higher resolution structures explain Lis1’s inability to bind dynein at two sites in the straight linker state

Previous low-resolution data showed that dynein in a strong microtubule binding state binds one Lis1 β-propeller at site ^40^. However, that structure was in a straight linker state, emphasizing the linker position as a critical aspect in Lis1 binding to dynein. Our straight linker map also had one Lis1 β-propeller bound (Fig. 3c). Aligning straight linker dynein with or without Lis1 by AAA1 shows that the linker shifts slightly to accommodate for Lis1 binding (Fig. 3f). We also saw clear density for the second Lis1 β-propeller in the intermediate state dynein, even though the nucleotide densities in this state are consistent with the ones in the straight linker dynein (Fig. 3b and Extended Fig. 4b and 6). Comparison of this state (I6, 3.4 Å resolution) with our straight linker dynein (I3, 4 Å resolution) bound to one Lis1 β-propeller showed that the site_ring_ interactions are conserved while site_stalk_ is shifted away from Lis1 as dynein goes from the intermediate to the straight linker state (∼15 Å) and is therefore not accessible for the second Lis1 β-propeller to bind (Fig. 3g and h). Binding of Lis1 at the second site is likely not to happen without the cooperativity from Lis1 bound at site_ring_. This is consistent with previous lower resolution structures suggesting that the straight linker dynein conformation is incompatible with the second Lis1 β-propeller binding^32,40,41^.

**Figure 4:**
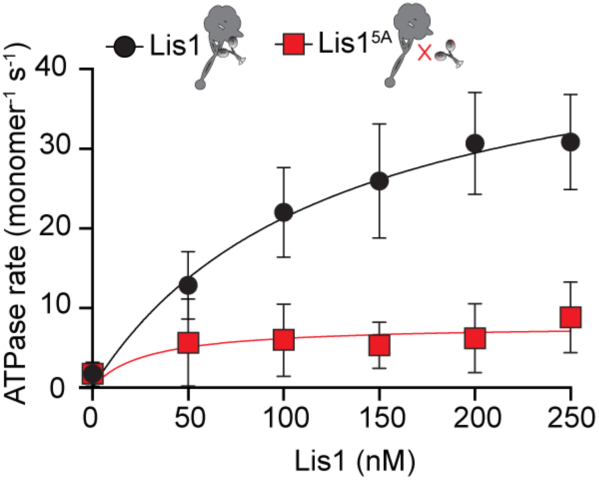
Lis1 increases dynein’s basal ATPase activity. ATPase activity of dynein in the presence of increasing concentrations of wild-type Lis1 (black dots) or a Lis1 mutant that does not bind to dynein (Lis1^5A^, red squares). Mean values (+/- SD) from three independent experiments are shown. Fitted values (+/- standard error of the fit) kbasal= 0.837 +/- 0.08 sec^-1^, kcat[Lis1]= 25.75 +/- 2.4 sec^-1^.

In our straight linker dynein bound to one Lis1 at site_ring_ the density for the AAA5 loop is missing, while it is present in the intermediate or bent linker structures bound to one or two Lis1 (Fig. 3h – top panel and Extended Data Fig. 7a). Importantly, mutation of this loop (N3475 and N3476) or the corresponding residues D253 and H254 in Lis1 disrupts dynein’s interaction with Lis1 and leads to binucleate phenotypes in yeast, suggesting that this interaction may only take place when dynein transitions from straight to intermediate or bent linker states^32^.

## Lis1 increases dynein’s basal ATP hydrolysis rate

Our analysis showed that dynein sampled more conformations at the shorter time point in the presence of Lis1 (Fig. 1g and h, and Extended Fig. 5c). We hypothesized that Lis1 could accomplish this by increasing dynein’s basal ATP hydrolysis rate. Several studies have shown that Lis1 does not affect dynein’s microtubule-stimulated ATPase activity^21,41,43^. However, the contribution of Lis1 to dynein’s basal ATPase hydrolysis rate (in the absence of microtubules) has not been fully established^21,41^. To test our hypothesis, we incubated the same dynein monomer (amino acids 1352- 4092) as the one used in cryo-EM studies with increasing concentrations of Lis1 and measured dynein’s basal ATP hydrolysis rate. We observed a Lis1 concentration-dependent increase in ATP hydrolysis rate (Fig. 4). Importantly, binding of Lis1 to dynein is required for this effect as the rate did not increase in the presence of a Lis1 mutant that does not bind to dynein (Lis1^5A^: R212A, R238A, R316A, W340A, K360A) (Fig. 4)^40,41,52^. This observation supports our hypothesis that Lis1 binding allows dynein to sample more conformations by increasing dynein’s basal ATP hydrolysis rate.

**Figure 5:**
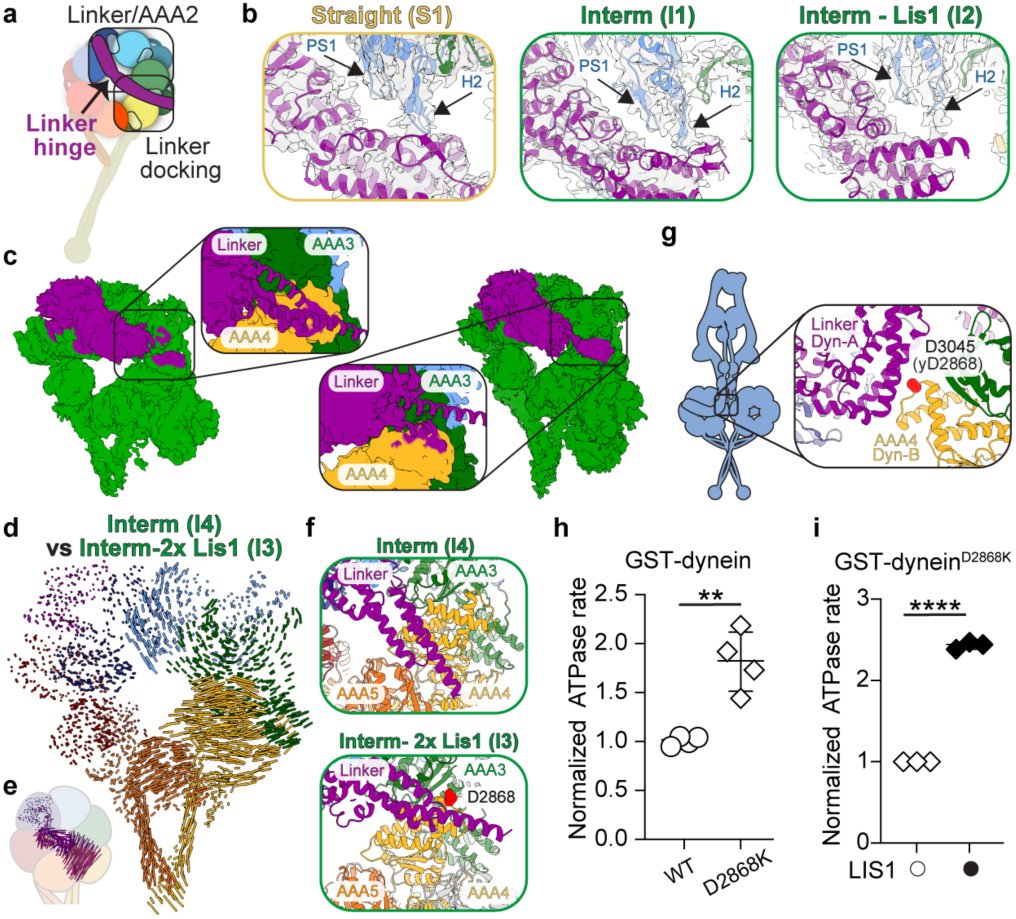
Conformational changes and Lis1 binding regulate dynein’s basal ATP-hydrolysis rate. **a**. Schematic of the two important interaction regions between dynein’s linker and motor domain. **b**. The proximity of the P1 sensor (PS1) and H2 insert loops in AAA2 (blue) to the linker (magenta) hinge region in the indicated models. **c**. Density maps and close-ups for subsets of particles selected from cryoDRGN analysis that showed the most extensive linker density. Close-ups show predicted linker (magenta) docking sites on AAA4 (yellow) after fitting an extended linker model in the density maps. **d**. Map of pairwise alpha-carbon distances between the intermediate linker model and the intermediate linker model bound to two Lis1 β-propellers. The models were aligned by their AAA1 modules. Residues 1361-1772 of the linker were removed for clarity. **e.** Residues 1352-1772 of the linker from the map of pairwise alpha-carbon distances between the intermediate linker model and the intermediate linker model bound to two Lis1 β-propellers in d. **f.** View of linker docking in intermediate linker dynein bound to Lis1 β-propellers (top panel) and intermediate linker dynein (bottom panel). Residue D2686 is highlighted in red. **g.** View of the location of residue D3045 (yeast D2868) in human dynein Phi particle (PDB: 5NVU). **h.** Normalized ATPase activity (median ± interquartile range) of GST-dynein and GST-dynein^D2868K^. **, p = 0.0022. Data from four independent experiments. **i.** Normalized ATPase activity (median ± interquartile range) of GST-dynein^D2868K^ in the absence (white circles) or presence (black circles) of Lis1. ****, p < 0.0001. Data from three independent experiments.

## Dynein activation is driven by conformational changes mediated by ATP-hydrolysis

To understand how Lis1 is increasing dynein’s ATPase rate, we further analyzed the interactions that the linker makes with the dynein motor domain, as these interactions have been previously shown to be important for dynein’s mechanochemistry^11,41,53^. We reasoned that the linker conformation in the different dynein states could explain how Lis1 regulates dynein’s mechanochemistry given the changes in linker position associated with ATP hydrolysis and Lis1 binding.

Two models for how swinging of the linker is regulated during ATP hydrolysis have been proposed. In the linker/AAA2 model, loops in AAA2 (pre-sensory I – PS1 and helix 2 – H2) rearrange during ATP hydrolysis in AAA1 leading to an interaction with the linker hinge and subsequent bending and swinging of the linker (Fig. 5a). Consistent with this, deleting the H2 region in *Dictyostelium* dynein reduces dynein’s basal and microtubule-stimulated ATPase activity^11^. In our straight and intermediate dynein state, when Lis1 is not bound to dynein the linker is contacting the AAA2 H2 loop, but these contact sites are less resolved when Lis1 is bound, suggesting lesser engagement of this region (Fig. 5b left and middle panels vs. right panel). This could mean that the need for this interaction is bypassed when Lis1 is bound to dynein due to the Lis1-driven increased flexibility in dynein.

In the linker docking model, ATPase-induced changes in AAA1 propagate to the linker docking interface on the motor domain to induce linker swing (Fig. 5a). Because our models lacked density for the linker docking sites when Lis1 was bound to dynein in the intermediate state, we performed heterogeneity analysis of all particles that belong to the intermediate state from the dataset collected in the presence of Lis1 and at the 0.5 min time point (Extended Data Fig. 2b and 4b – I1). In addition to separating the populations that have one or two Lis1 β-propellers bound, we observed other conformational changes that correspond to linker movement (Extended Data Fig. 2a). We then picked particles from the states that showed the most linker density, regardless of whether they had one or two Lis1 β-propellers bound. Although this analysis did not improve the resolution of our maps, it did allow us to extend the modeling of the linker in the region where it could potentially dock on dynein. For each density map generated in this analysis, the linker comes close to AAA4 or the interface between AAA3 and AAA4 (Fig. 5c and Extended Data Fig. 2c).

We next compared these linker docking interfaces with intermediate-state dynein observed in the absence of Lis1. The presence of Lis1 led to conformational changes in AAA1 through AAA5 (Fig. 5d). The linker is also shifted, with the docking site much closer to the interface between AAA4 and AAA5 when Lis1 is not present (Fig. 5e and f). Interestingly, in the Lis1-bound dynein, the linker is close to D2868 (3045 in human dynein), a residue that makes important contacts with the linker of the opposing dynein in the Phi particle (Fig 5f – bottom panel and g)^5^. A charge-reversing mutation in this residue disrupts Phi particle formation and almost completely bypasses the need for Lis1 in yeast^5,38^. Given that Phi traps dynein in a single conformation, and that conformational changes such as opening and closing of the ring and linker swing are required for dynein to go through its mechanochemical cycle, we reasoned that the Phi conformation inhibits dynein’s mechanochemistry, in addition to preventing it from binding to microtubules. This leads to the prediction that disrupting Phi particle formation would increase dynein’s basal ATPase hydrolysis rate. To test this, we used GST-dynein, as the Phi-disrupting D2868K mutation in this construct does not affect dynein’s motile properties or Lis1 binding^38^. The ATPase rate of GST-dynein-D2868K was almost double that of the wild-type construct, suggesting that Phi conformation inhibits dynein’s ability to hydrolyze ATP (Fig. 5h). Addition of Lis1 continued to increase dynein’s ATPase rate implying that there are likely other interactions important to this process.

## Lis1 binding to dynein might be facilitating nucleotide release from AAA3

Finally, to better understand Lis1’s effects on dynein’s mechanochemistry, we turned to all-atom molecular dynamic (MD) simulations. We first asked how the presence of Lis1 influences dynein’s linker bending, given the importance of linker position in the mechanochemical cycle (Fig. 5a)^11,53^. To do this, we first defined the linker rotation angle between residues S1942 (AAA1), L1664 (linker), and K1424 (extended part of the linker) in our highest resolution dynein models obtained in this work: bent-1x Lis1 (Fig. 3e – B7), interm-1x Lis1 (Fig. 5d – I6), interm-2x Lis1 (Fig. 5d – I7), and interm dynein (Fig. 2b – I4), and determined from heterogeneity analysis, in which we modelled the same number of residues for the linker region (Methods and Fig. 6a). For each state, we launched multiple Gaussian Accelerated MD simulations (Methods), which can enhance the conformational sampling of molecular machines, and measured population density for the different linker angles. Our analysis shows that the linker rotation is more constrained when Lis1 is bound to dynein in the intermediate state (Fig. 6b). Likely, this is caused by the steric hindrance from Lis1 binding do dynein.

**Figure 6:**
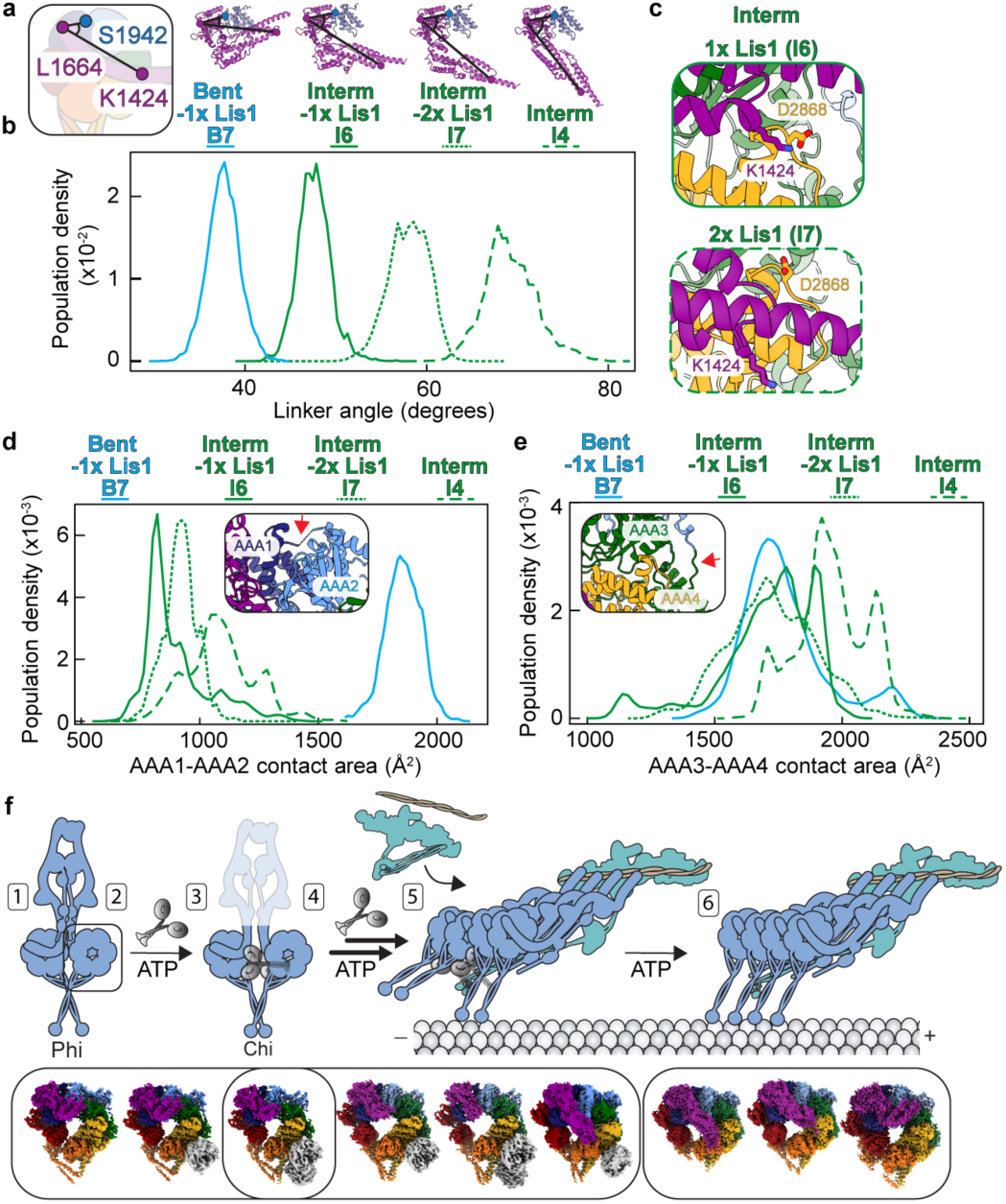
Molecular dynamics support changes in dynein conformations due to Lis1 binding. **a**. The linker rotation angles defined by the Cα atoms of residues located in AAA2 (S1942, corner) and linker – corner (L1664 – S1924, edge 1) and linker – corner (L1664 – K1424, edge 2) for the four systems used in the molecular dynamic simulations: bent linker with one Lis1 β-propellers bound (B7, solid blue line), intermediate linker with one Lis1 β-propeller bound (I6, solid green line), intermediate linker with two Lis1 β- propellers bound (I7, dotted green line), and intermediate linker (I4, large dot green line). **b.** Population density for the linker rotation angles for the four simulation systems. **c.** Predicted salt bridge between D2868 in AAA3 and K1424 in the linker for intermediate linker with one Lis1 β- propeller bound (I6 – top panel) and intermediate linker with two Lis1 β- propellers bound (I7 – bottom panel). **d.** Distribution of contact area surface between AAA1 and AAA2 (insert – red arrow points to the interface) in the simulations. **e.** Distribution of contact area surface between AAA3 and AAA4 domains (insert – red arrow points to the interface) in the simulations. **f.** Model of dynein activation and the proposed placement of the identified structures in the model. 1 – dynein in the Phi particle, 2 – initial step of Chi particle formation, 3 – Chi particle, 4 – increase in dynein’s basal ATP hydrolysis rate, 5 – full dynein complex assembles with dynactin and an activating adaptor, and binds to microtubules, 6 – Lis1 dissociates from active dynein complex.

We next evaluated if the residue D2868, a key residue in maintaining Phi particle formation could be making contacts with the linker when Lis1 is bound to dynein. We found that D2868 forms a salt bridge with K1424 (linker), which could stabilize the intermediate linker position when Lis1 is bound to dynein in this state (Fig. 6c). This salt bridge is relatively stable in the simulation with the intermediate dynein bound to one Lis1 β-propeller (Fig. 6c – top panel), whereas a disengaged configuration is preferred in the intermediate dynein bound to two β-propellers (Fig. 6c – bottom panel).

Lastly, we asked whether the presence of Lis1 could be affecting the nucleotide binding pockets in AAA1 and AAA3 in our simulations. The analysis of the contact area between AAA1 and AAA2 revealed that the bent linker enables a tighter interface than that in the intermediate linker state, presumably allowing this site to become more hydrolysis competent (Fig. 6d). This is consistent with closing of the AAA1 interface in the bent conformation seen for *Dictyostelium* dynein in the pre-power stroke state primed for ATP hydrolysis^48^. We next investigated the AAA3 and AAA4 contact area and showed that Lis1 binding loosens the AAA3 and AAA4 interface, potentially favoring ADP release from the AAA3 binding pocket (Fig. 6e). Interestingly, binding of the second Lis1 β-propeller further loosens the interface, as illustrated by comparing the population distributions of interm-2x Lis1 vs. interm-1x Lis1 systems (Fig. 6e, dotted vs. solid green lines). This analysis suggests that Lis1 might be increasing dynein’s basal ATP hydrolysis rate by facilitating nucleotide release from AAA3.

Our work showed how heterogeneity mining in cryo-EM data combined with time-resolved approaches could identify a conformational landscape for dynein during its mechanochemical cycle and how this landscape is altered by Lis1 binding. The conformations identified in our study are in line with a recent similar analysis of full-length human dynein^51^. However, our work goes further, defining the contributions of Lis1 to dynein’s mechanochemistry. Previous attempts at capturing different steps of dynein’s mechanochemical cycle using X-ray crystallography were not fully successful. Combining heterogeneity mining with time-resolved cryo-EM provides a unique advantage to visualize how ATP hydrolysis drives changes in dynein’s conformational landscape over time.

We identified multiple conformations of dynein during ATP hydrolysis and determined the nucleotide state of each AAA module. Our different dynein and dynein/ Lis1 structures can be mapped onto the dynein activation pathway, starting with the bent linker dynein without Lis1. This conformation is consistent with individual dynein protomers in Phi (Fig. 6f-1). The bent linker dynein with one Lis1 β-propeller bound at site_ring_ likely represents an early step in Chi particle formation (Fig. 6f-2), followed by binding of the second Lis1 β-propeller at site_stalk_ and full assembly of Chi (Fig. 6f-3). We also identified a small population of particles in our datasets that corresponded to Chi dynein, containing two dyneins and two Lis1 dimers (published previously in^36^). Thus, our analysis is consistent with the idea that Lis1 binding to dynein and formation of Chi is an early step in dynein activation.

We propose a model in which Lis1 binding to dynein initially disrupts the autoinhibited Phi conformation by forming the Chi particle (Fig. 6f-3)^36^. Once bound, Lis1 allows dynein to go through its mechanochemical cycle, which is seen as an increase in dynein’s basal ATP hydrolysis rate (Fig. 6f-4). This likely both disrupts Chi and allows dynein to adopt conformations compatible with complex assembly with dynactin and an activating adaptor (Fig. 6f-5). Interestingly, we did not observe intermediate linker dynein by itself in any of our datasets collected in the presence of Lis1, even though we could solve structures of straight and bent linker dynein with and without Lis1 bound from those same datasets (Fig. 1g and Extended Data Fig. 4a and b). In addition, at the 30 min timepoint and in the presence of Lis1, the intermediate state dynein was the dominant one (Fig. 1h and Extended Data Fig. 5c). Although we cannot rule out that sample preparation somehow led to these distributions, this could mean that binding of Lis1 to dynein is particularly favorable in the intermediate state. Once the initial complex assembles and the first set of dynein dimers binds to microtubules (Fig. 6f-5), Lis1 may dissociate from dynein to allow for dynein’s movement on microtubules (Fig. 6f-6). An alternative hypothesis suggesting the presence of Lis1 moving with active dynein is also possible given recent cellular work showing colocalization of moving dynein and Lis1 in neuronal cells^54^. The dynein conformations we observed in the absence of Lis1 likely represent the different steps of dynein’s ATP hydrolysis-driven movement on microtubules. Further structural work with full-length dynein, dynactin, activating adaptor, and Lis1 in the presence of ATP will be required to fully map dynein’s release from autoinhibition and activation.

We showed direct evidence that Lis1 stimulates dynein’s basal ATP hydrolysis rate and suggest that this basal activity is inhibited when dynein is in the Phi conformation. In accordance with this, the D2868K mutation in dynein that prevents Phi formation increases dynein’s basal ATPase rate. However, Lis1 stimulates dynein’s ATPase activity further after the Phi particle is disrupted. This suggests a two-step regulation mechanism. In the first step, Lis1 binds to dynein to open the Phi particle, which results in a higher rate of ATP hydrolysis as the conformational constraints imposed by Phi are released. Once Phi has been disrupted, Lis1 further increases dynein’s ATPase rate by acting on the released motor, possibly by facilitating the release of ADP from AAA3. This could be driven by the decreased communication between the linker and dynein’s motor domains, as suggested by the MD simulations. There, Lis1 binding to dynein in the intermediate state appeared to be influencing the ability of the linker to communicate with AAA4 (D2868) and the H2 loops in AAA2 (Fig. 5b). Further structural and mechanistic work will be required to fully decipher this mechanism.

## Acknowledgements

We thank the cryo-EM Facility at UC San Diego. We thank Janice Reimer for helpful discussions. We also thank our funding sources: AAK was supported by American Cancer Society PF-18-190-01-CCG; KHVN is supported by the Molecular Biophysics Training Grant (NIH grant T32 GM139795); WM is funded by an Early Career Research Award from the Cardiovascular Research Institute of Vermont; EPK is funded by a Jane Coffin Childs Postdoctoral Fellowship; AL’s lab by NIH R01 GM107214 and R35 GM145296; and SRP’s lab by the Howard Hughes Medical Institute and NIH R35 GM141825. This paper was typeset with the bioRxiv word template by @Chrelli: www.github.com/chrelli/bioRxiv-word-template.

## Data availability

Cryo-EM maps and atomic coordinates have been deposited in the Electron Microscopy Data Bank under accession codes: S1 – EMD-46919, S2 – EMD- 46897, S3 – EMD-46938, S4 – EMD-46953, I1 – EMD-47019, I2 – EMD-46958, I3 – EMD-46941, I4 – EMD-46954, I5 – EMD-46974, I6 – EMD-46972, I7 – EMD-46975, B1 – EMD-46935, B2 – EMD-46940, B3 – EMD-46942, B4 – EMD-46919, B5 – EMD-46959, B6 – EMD-47033, B7 – EMD-47026, B8 – EMD-47032 and in the Protein Data Bank under accession codes: S1 – 9DIU, S2 – 9DI3, S3 – 9DJU, S4 – 9DKD, I1 – 9DMW, I2 – 9DKH, I3 – 9DJZ, I4 – 9DKE, I5 – 9DLD, I6 – 9DKX, I7 – 9DLE, B1 – 9DKM, B2 – 9DJY, B3 – 9DK0, B4 – 9DJ7, B5 – 9DKJ, B6 – 9DNB, B7 – 9DN5, B8 – 9DN7.

## Materials and Methods

### Yeast Strains

The *S. cerevisiae* strains used in this study are listed in Extended Data Table 1. The endogenous genomic copy of *PAC1* (encoding Lis1) or *DYN1* (encoding dynein) were deleted using PCR-based methods as previously described^55^. Point mutation to generate the D868Q dynein was introduced using QuikChange site-directed mutagenesis (Agilent) with the primers: 5’- CTTTAGGTCTTTTATTGAAGACAGAACAAGAACTG-3’ and 5’-CAG- TTCTTGTTCTGTCTTCAATAAAAGACCTAAAG-3’ and verified by DNA sequencing. The mutated fragment was re-inserted into the kl*URA3* strains using the lithium acetate method^56^ to reintroduce the mutated *DYN1* gene. Positive clones (kl*URA3-*) were selected in the presence of 5-Fluorootic acid, screened by colony PCR, and verified by DNA sequencing.

### Protein expression and purifications

Protein purification steps were done at 4°C unless otherwise indicated. *S. cerevisiae* dynein constructs were purified from *S. cerevisiae* using a ZZ tag as previously described ^57^. Briefly, liquid nitrogen-frozen yeast cell pellets were lysed by grinding in a chilled coffee grinder and resuspended in Dynein-lysis buffer supplemented 0.5 mM Pefabloc, 0.05% Triton, and cOmplete EDTA-free protease inhibitor cocktail tablet (Roche). The lysate was clarified by centrifuging at 264,900 × *g* for 1 hr. The clarified supernatant was incubated with IgG Sepharose beads (GE Healthcare Life Sciences) for 1 hr. The beads were transferred to a gravity flow column, washed with Dynein-lysis buffer supplemented with 250 mM potassium chloride, 0.5 mM Pefabloc and 0.1% Triton, and with TEV buffer (10 mM Tris–HCl [pH 8.0], 150 mM potassium chloride, 10% glycerol, and 1 mM DTT). Dynein was cleaved from IgG beads via incubation with 0.15 mg/mL TEV protease (purified in house) overnight at 4°C. Cleaved dynein was concentrated using 100K MWCO concentrator (EMD Millipore), filtered by centrifuging with Ultrafree-MC VV filter (EMD Millipore) in a tabletop centrifuge and used fresh for cryo-EM sample preparation or snap-frozen in liquid nitrogen.

Yeast Lis1 was purified from *S. cerevisiae* using 8xHis and ZZ tags as previously described^21^. In brief, liquid-nitrogen frozen pellets were ground in a pre-chilled coffee grinder, resuspended in buffer A (50 mM potassium phosphate [pH 8.0], 150 mM potassium acetate, 150 mM sodium chloride, 2 mM magnesium acetate, 5 mM β-mercaptoethanol, 10% glycerol, 0.2% Triton, 0.5 mM Pefabloc) supplemented with 10 mM imidazole (pH 8.0) and cOmplete EDTA-free protease inhibitor cocktail tablet, and spun at 118,300 x g for 1h. The clarified supernatant was incubated with Ni-NTA agarose (QIAGEN) for 1 hr. The Ni beads were transferred to a gravity column, washed with buffer A + 20 mM imidazole (pH 8.0), and eluted with buffer A + 250 mM imidazole (pH 8.0). The eluted protein was incubated with IgG Sepharose beads for 1 hr. IgG beads were transferred to a gravity flow column, washed with buffer A + 20 mM imidazole (pH 8.0) and with modified TEV buffer (50 mM Tris–HCl [pH 8.0], 150 mM potassium acetate, 2 mM magnesium acetate, 1 mM EGTA, 10% glycerol, 1 mM DTT). Lis1 was cleaved from the IgG beads by the addition of 0.15 mg/mL TEV protease (purified in house) for 1 hr at 16°C. Cleaved proteins were filtered by centrifuging with Ultrafree-MC VV filter (EMD Millipore) in a tabletop centrifuge and flash frozen in liquid nitrogen.

### Electron microscopy sample preparation

To remove any residual nucleotide, purified fresh dynein (not frozen) was incubated with apyrase for 30 mins (apyrase, 0.1 U/ml, NEB) prior to gel filtration on a Superose 6 Increase column pre-equilibrated in buffer containing 50 mM Tris, pH 8, 150 mM KCl, 2 mM EDTA, and 1 mM DTT. Eluted protein was concentrated to about 5 µM and diluted 1:1 with Lis1 or TEV buffer and incubated on ice for 10 mins prior to ATP addition (final concentrations: 2.5 µM dynein, 2.5 µM Lis1, and 1.25 mM ATP). Samples containing ATP were incubated for a total of 0.5 min or 30 min on ice (including blotting time and plunging) and quickly applied to plasma cleaned (Solarus, Gatan) UltrAuFoil Holey Gold R 1.2/1. µM3 grids (Quantifoil). A vitrobot (FEI) was used to blot away excess sample and plunge freeze the grids in liquid ethane. The Vitrobot chamber was maintained at 100% humidity and 4°C during the process. Grids were stored in liquid nitrogen until ready to be imaged. The 0.5 min grids were made and analyzed first. Based on observed conformational landscape in this condition, the 30 min time point was chosen for the longer time condition.

### Electron microscopy image collection

The 0.5 min time point grids were imaged using Talos Arctica operated at 200 kV and equipped with K2 Summit direct electron detector (Gatan). Automated data collection was performed using Leginon. A total of 1849 movies over 3 sessions for the dynein + ATP – 0.5-min dataset and 12370 movies over 4 sessions for the dynein + Lis1 + ATP – 0.5-min dataset at 36,000 magnification (1.16 Å/pixel). The dose was ∼6.3 Å^-2^s^-^1 with a total exposure time of 11 s divided into 200 ms-frames for a total of 40 frames. The defocus range was set to 0.8 – 2.4 µm.

The 30 min time point grids were imaged using an FEI Titan Krios (200 kV) equipped with a Flacon 4i detector and energy filter (<10 eV slit size) (Thermo Fisher). Total of 7182 movies for the dynein + ATP – 1800 sec dataset and 6670 movies for the dynein + Lis1 + ATP – 1800 sec dataset were collected using automated data acquisition (Thermo Fisher EPU) at 130,000 magnification (0.889 Å/pixel). The dose was ∼5 Å^-2^s^-^1 with a total exposure time of 10 s divided into 200 ms-frames for a total of 40 frames. The defocus range was set to 0.75 – 2.8 µm.

### Electron microscopy data processing

All movies were aligned in CryoSPRC live^58^ using MotionCor2^59^ of the dose-weighted frames. CTF was estimated in dose-weighted images using CTFFIND4^60^. Images with CTF fits worse than 5 Å were excluded from further processing. Particles were initially selected using blob finder in CryoSPRC and these peaks were used for Topaz^61^ model training and final particle picking using Topaz. Particles were extracted and binned to 4.64 Å/pixel for the 0.5 min dataset and to 3.556 Å/pixel for the 30 min dataset. Multiple rounds of 2D classification with a varying number of online-EM interactions (30-40) and batch size per class between 200-400 dependent on the number of particles in each classification were carried out first in cryoSPARC to remove bad particles. Reconstructions were performed to generate initial models for each dataset with particles from all good 2D classes. Good particles and models were selected for multiple rounds of 3D refinement and local refinements until no further improvement was observed and models appeared to contain well-defined secondary structures. Further analysis of heterogeneity was performed using 3D classification without alignment in Relion-3 to find subsets of particles that classified into models with well-defined secondary structures. These particles were then transferred to cryoSPARC for another round of Ab-initio reconstructions and 3D refinement.

### CryoDRGN classification

To identify conformational and compositional variability, volumes from the non-uniform refinement in cryoSPARC were further subjected to cryoDRGN^62,63^. First, the particles were binned from 352 pixels to 128 pixels (3.19 or 2.45 Å/pixel final pixel size) and used for training an 8-dimensional latent variable model with 3 hidden layers and 128 nodes in the encoder and decoder networks. The latent space was visualized with the analyze function on epoch 50 and clusters were extracted using k-means analysis. The particles from the cluster that yielded the best 3D refinement were moved forward for one more round of training, with binning to 256 pixels (1.595 or 1.22 Å/pixel final pixel size) and training with 3 hidden layers and 1024 nodes in the networks. cryoDRGN landscape analysis was performed on the latent space and the structures representing the center of each cluster were manually inspected. Finally, good particles selected in cryoDRGN were extracted and transferred cryoSPARC for another round of Ab-initio reconstructions and 3D refinement.

### Model building and refinement

Local refinement, CTF, and de-focus optimizations were used to improve the overall resolution of each map. The individual domains (AAA, AAA2, AAA3, AAA4, linker, and different stalk helices) from yeast dynein models PDB: 7MGM and 5VH9, were docked, and rigid body fit into the different cryo-EM maps using UCSF Chimera X to build uniform models^64^. Subsequent rigid body and refinement was carried out using Phenix real space refine^65^. For all models the discrepancies between the model and map were fixed manually in COOT and then refined using a combination of Phenix real space refine and Rosetta Relax (v.13)^66,67^. For the 0.5 min dataset collected in the absence of Lis1 one model was built for the highest resolution density map (S1), given that the additional straight and intermediate state maps were derived from particles in S1 and were at a much lower resolution.

### ATPase assays

ATPase assays were performed using EnzChek phosphate kit (Thermo Fisher) as previously described^57^. The proteins for the final reaction mixtures contained 20 nM dynein monomer or 20 nM dynein GST-dimer and Lis1 0-250 nM, 2 mM Mg-ATP, 200 mM MESG (2-amino-6-mercapto-7methyl purine ribose), 1 U/ml purine nucleoside phosphorylate, and assay buffer (30 mM HEPES pH 7.4, 50 mM KOAc, 2 mM MgOAc, 1 mM EGTA, 1 mM DTT). The final reaction signal was read at 360 nm every 10 sec for 10 min on a Biotek Citation 5 plate reader.

### Molecular dynamics simulations

Four simulation systems were built based on the highest-resolution cryoEM models obtained at 30-min time point (bent linker with 1x Lis1 bound – B7, intermediate linker with 2x Lis1 bound – I7, intermediate linker with 1x Lis1 bound – I6, and intermediate linker with no Lis1 bound – I4). The linker region for each model was extended to start at residue 1354. For the intermediate linker dynein with 1x Lis1 bound – I6 and 2x Lis1 bound – I7, the linker position was determined from maps obtained from cryoDRGN heterogeneity analysis (Fig. 5c and Extended Data Fig. 2c; left panel for I6 and right panel for I7). Each model was solvated in a water box containing 150 mM NaCl. All the MD simulations were performed using the GPU-accelerated version of Amber18 with the ff14SB force field^68,69^. Energy minimization was conducted with the protein atoms’ positions constrained by harmonic potentials with a spring constant of 10 kcal/(mol Å^2^). With the same positional constraints, a following 40-ns equilibration simulation was performed at 300 K, gradually reducing the spring constant from 10 to 0.01 kcal/(mol Å^2^). Langevin dynamics with a friction coefficient of 1 ps^−1^ was applied to maintain a constant temperature. Particle Mesh Ewald was used for full-system periodic electrostatics while a 9 Å cutoff was applied to Lennard-Jones interactions^70^. Bonds involving hydrogen atoms were constrained using the SHAKE algorithm^71^. Next, Gaussian Accelerated Molecular Dynamics (GaMD) was employed to enhance the sampling of protein conformational dynamics^72^. For each system, 20 independent runs were launched, each comprising a 40-ns conventional MD stage, a 20-ns GaMD equilibration stage, and a 100-ns GaMD production stage. The conventional MD stage was used to gather statistics for calculating initial GaMD acceleration parameters. During the GaMD stages, both total potential energy boost and dihedral energy boost were applied to the system, each with an upper limit of 4 kcal/mol for the standard deviation (for accurate reweighting). For each system, the accumulated GaMD trajectories analyzed totaled 2µs. The probability density distributions shown in Fig. 6 were obtained using the reweighting approach^73^.

### Statistical Analysis

All statistical tests were generated using GraphPad Prism 9. The exact value of n, evaluation of statistical significance, P values, and specific statistical analysis are described in the corresponding figures and figure legends.

---

**Supplementary Video 1. Dynein conformations in the dynein/ Lis1 sample incubated with ATP for 0.5 min.**

The video shows a morph between different cryo-EM densities identified in the dynein and Lis1 dataset incubated for 0.5 mins with ATP.

**Extended Data Fig. 1.**
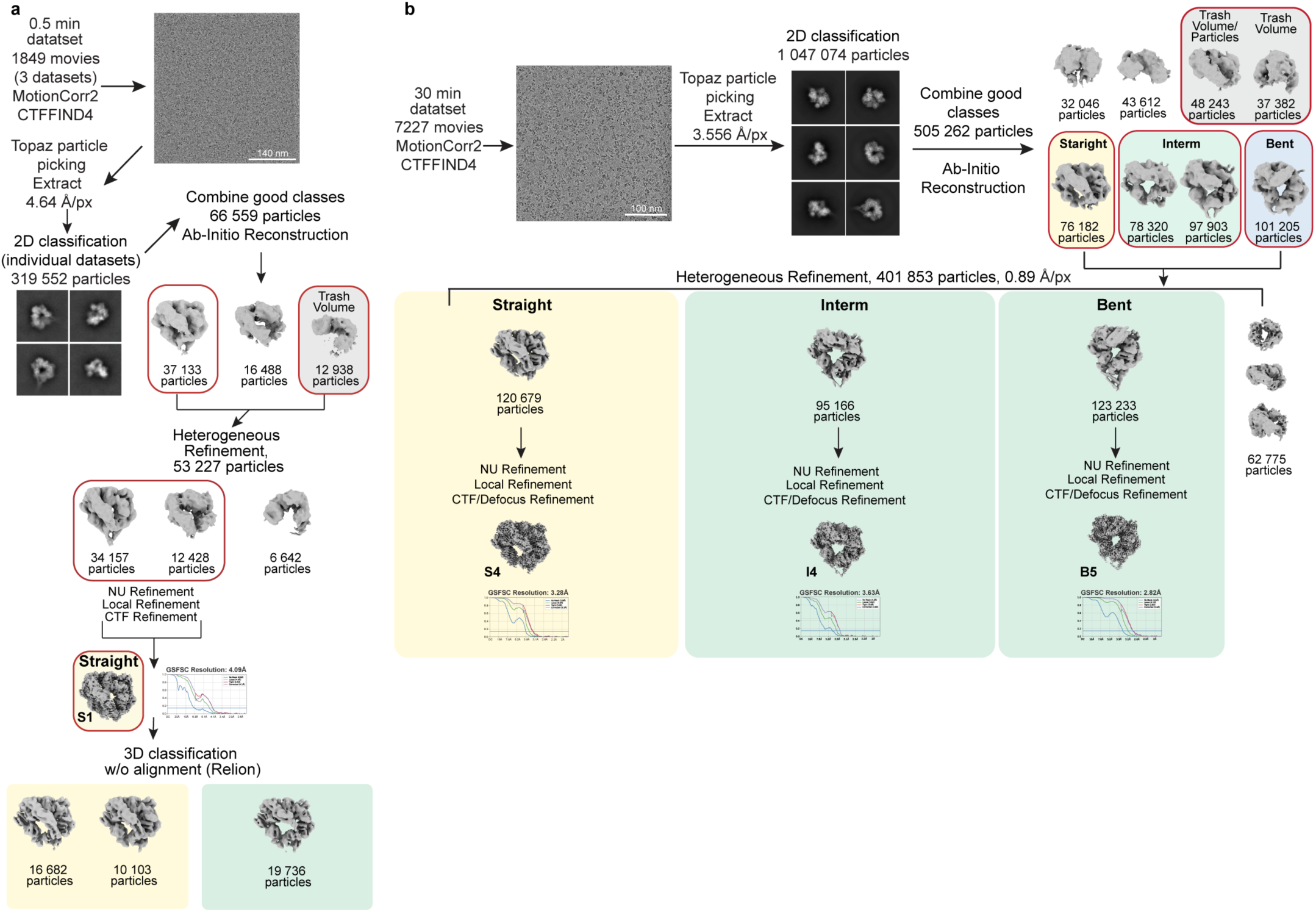
Cryo-EM data processing workflow for the dynein + Lis1 datasets. **a.** Dose-weighted movies from 3 datasets were aligned with MotionCor2 and CTF was estimated using CCTFFIND4. Particle extraction (binned by 4) was performed in cryoSPARC with a Topaz-trained model. Good particles from 2D classification jobs were used for ab-inito model generation in cryoSPARC. The models in red boxes were carried into the next steps. Fourier shell correlation (FSC) plots are shown next to the final maps. **b.** Dose-weighted movies from a dataset were aligned with MotionCor2 and CTF was estimated using CCTFFIND4. Particle extraction (binned by 4) was performed in cryoSPARC with a Topaz-trained model. Good particles from 2D classification jobs were used for ab-inito model generation in cryoSPARC. The models in red boxes were carried into the next steps. Fourier shell correlation (FSC) plots are shown next to the final maps.

**Extended Data Fig. 2.**
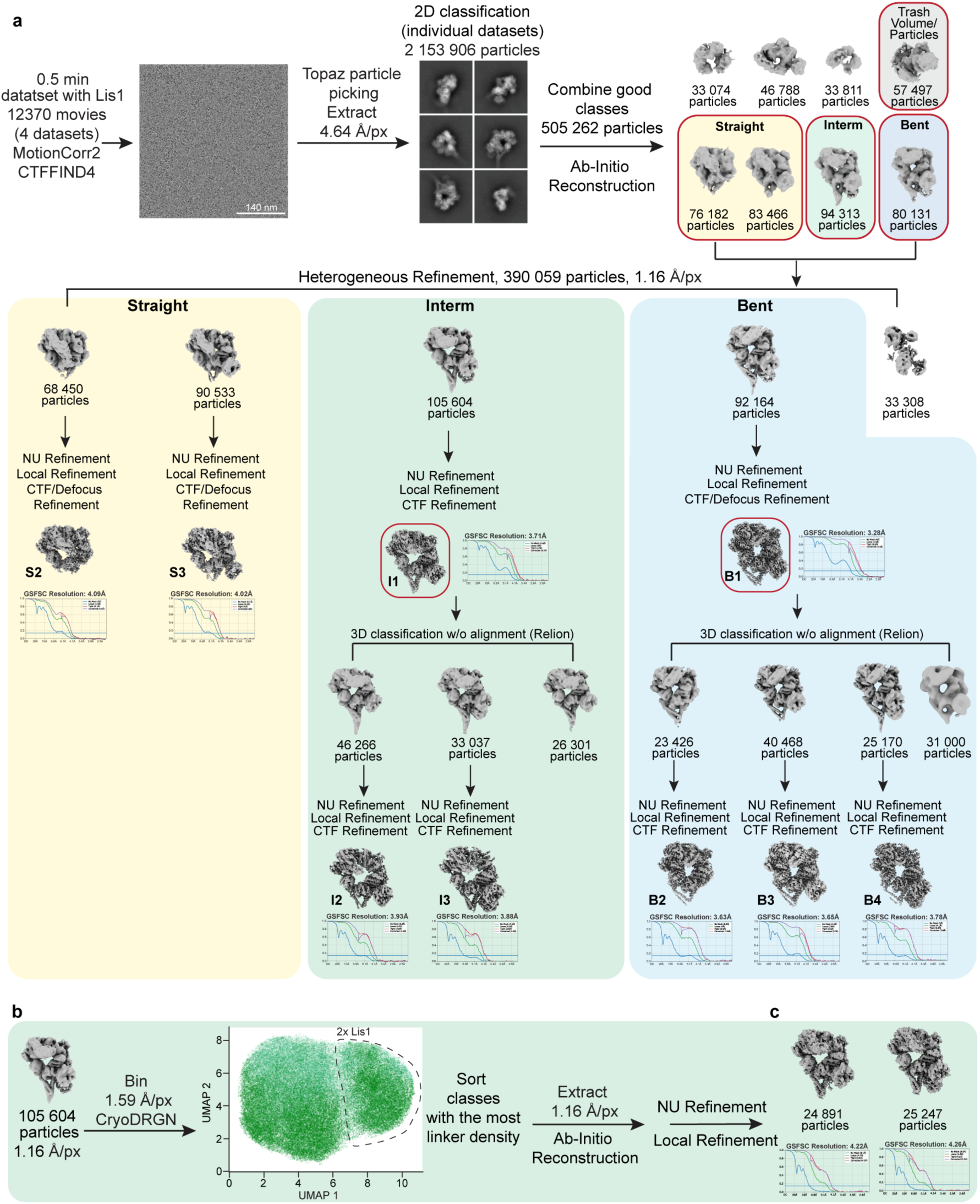
Cryo-EM data processing workflow for the dynein + Lis1 + ATP 0.5 min dataset. **a.** Dose-weighted movies from 4 datasets were aligned with MotionCor2 and CTF was estimated using CCTFFIND4 Particle extraction (binned by 4) was performed in cryoSPARC with a Topaz-trained model. Good particles from 2D classification jobs were used for ab-inito model generation in cryoSPARC. The models in red boxes were carried into the next steps. Fourier shell correlation (FSC) plots are shown next to the final maps. **b**. The heterogeneity analysis (cryoDRGN) workflow with particles belonging to the intermediate linker dynein (I1) conformation. A UMAP visualization after analysis is shown and particles that show binding of two Lis1 β-propellers are grouped into the highlighted population (2x Lis1). **c.** The two maps used to determine the linker position for the intermediate linker dynein with 1x Lis1 bound (left) and 2x Lis1 bound (right).

**Extended Data Fig. 3.**
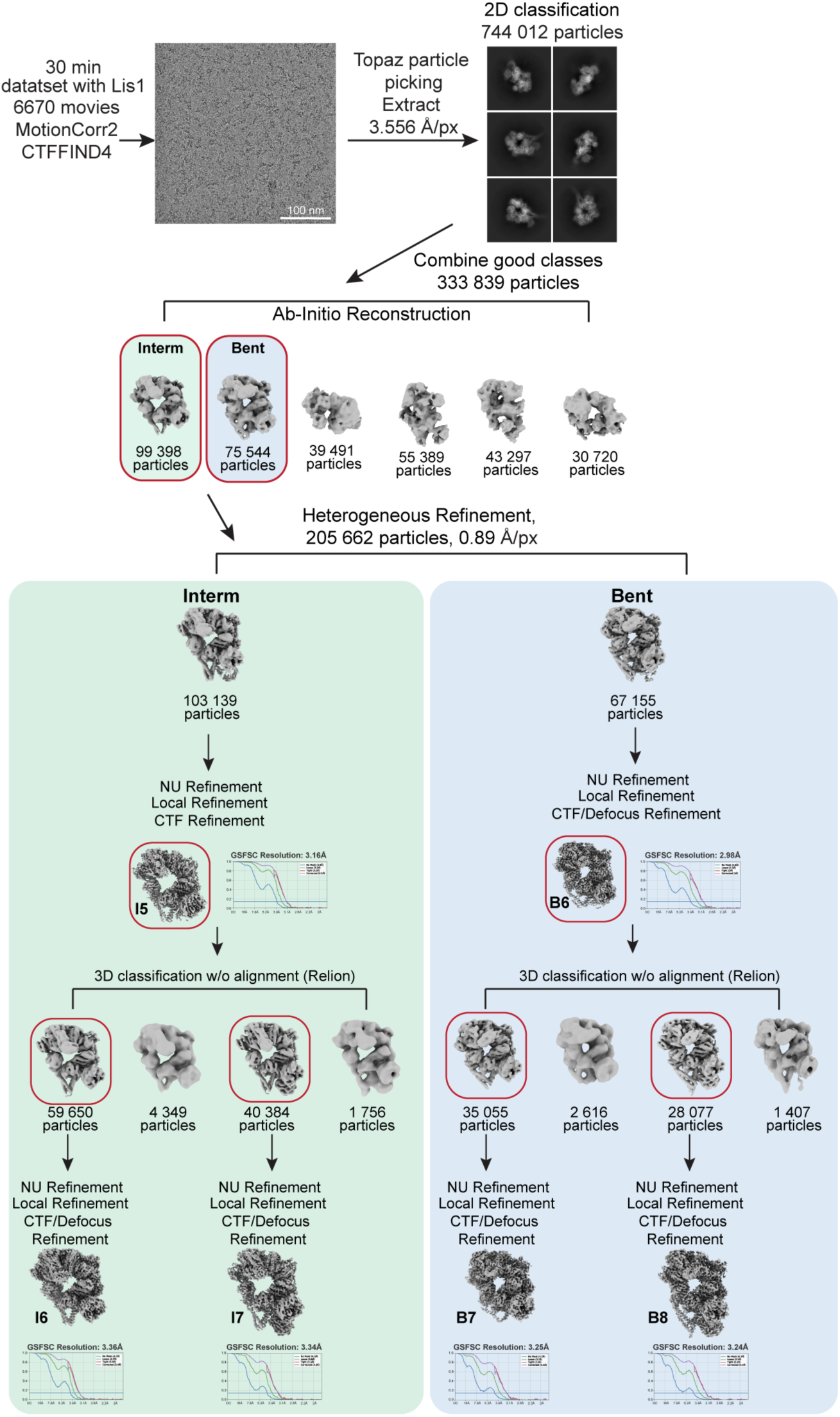
Cryo-EM data processing workflow for the dynein + Lis1 + ATP 30 min dataset. Dose-weighted movies from a dataset were aligned with MotionCor2 and CTF was estimated using CCTFFIND4. Particle extraction (binned by 4) was performed in cryoSPARC with a Topaz-trained model. Good particles from 2D classification jobs were used for ab-inito model generation in cryoSPARC. The models in red boxes were carried into the next steps. Fourier shell correlation (FSC) plots are shown next to the final maps.

**Extended Data Fig. 4.**
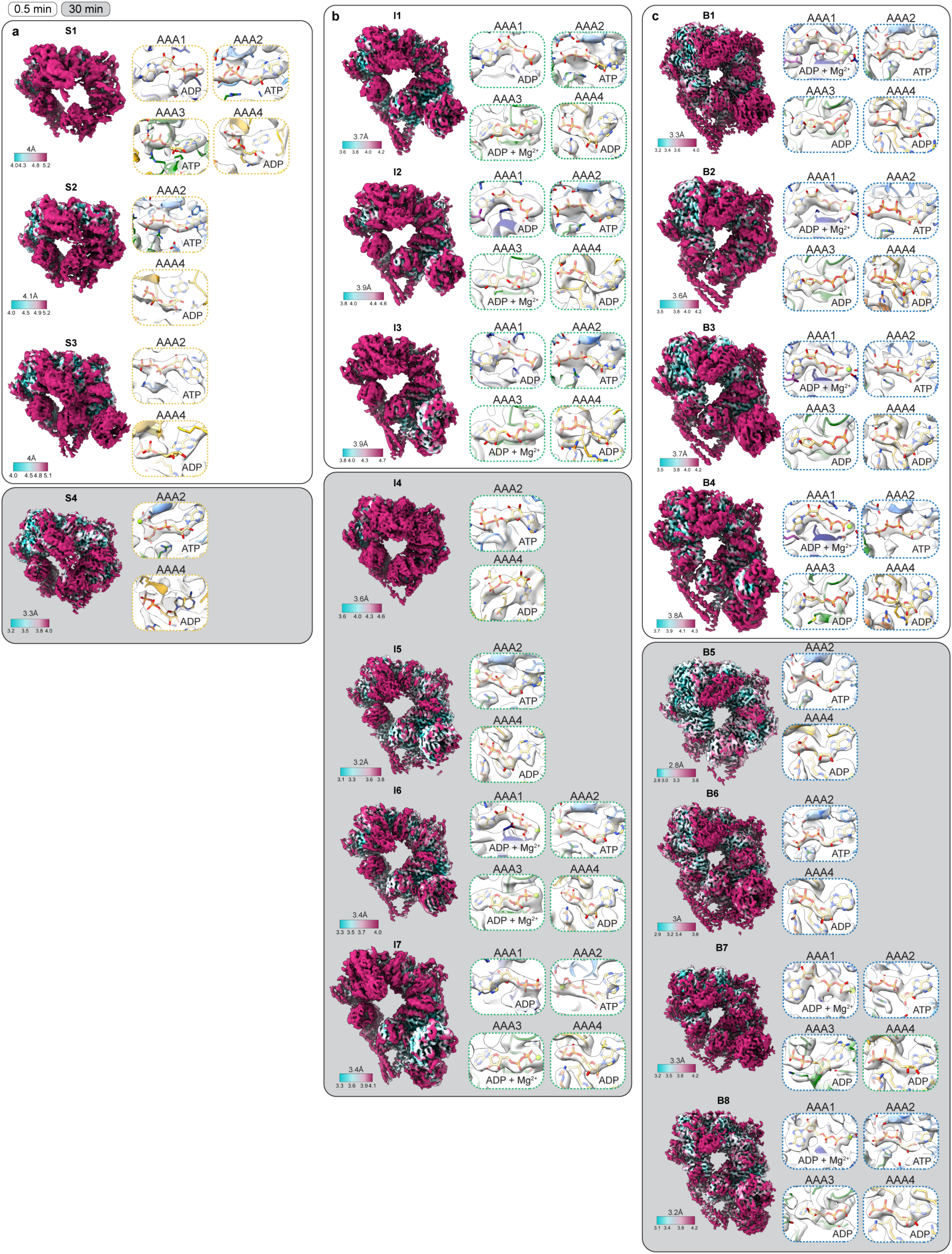
Local resolution and nucleotide occupancies at AAA+ subunits. Local resolution and views of the nucleotide-binding pockets for the indicated AAA+ subunits for **a**. straight (S), **b**. intermediate (I), and **c**. bent (B) linker dynein. Conformations belonging to each category are named with the first letter of that category and are shown in order starting with states identified in the shorter time point (0.5 min, white background) followed by the longer time point (30 min, gray background).

**Extended Data Fig. 5.**
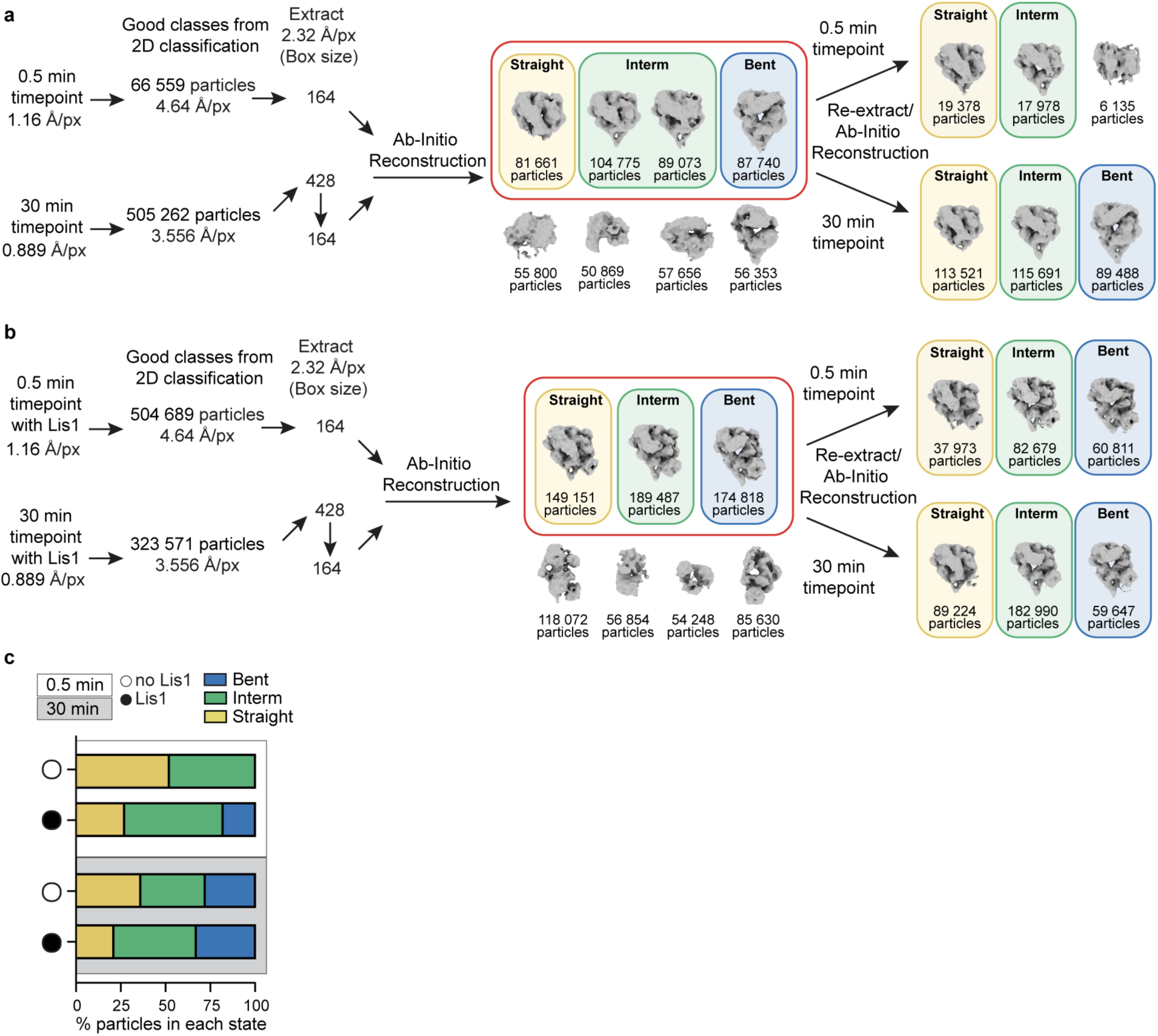
Cryo-EM data processing of the combined datasets. **a.** and **b.** Cryo-EM data processing workflow for the combined datasets. **c.** Relative abundance of particles belonging to different states obtained from particle distributions in cryo-EM datasets in the absence (open circle) or presence (black circle) of Lis1 at two different time points (0.5 min – white background and 30 min – gray background).

**Extended Data Fig. 6.**
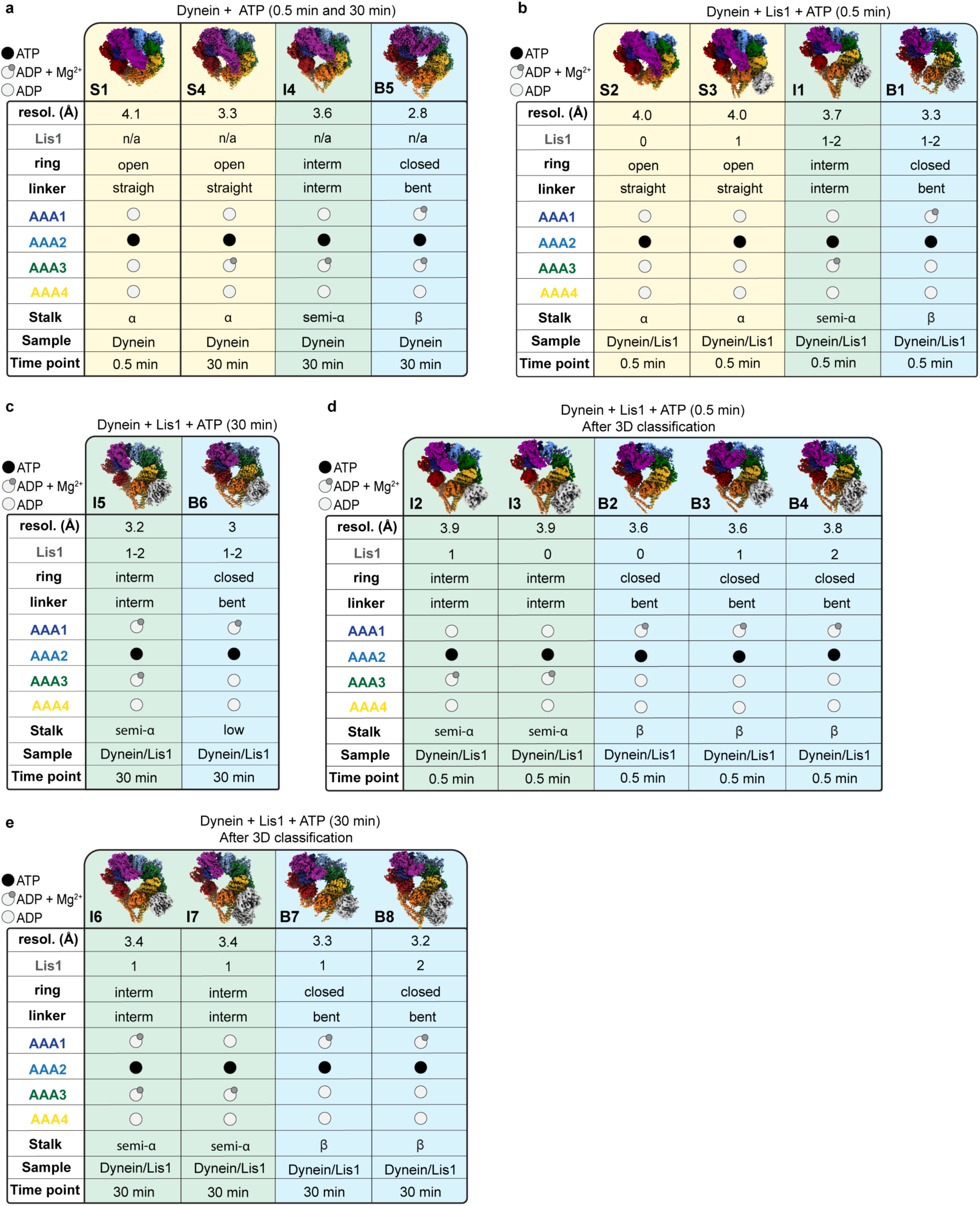
Summary of the identified structures and their properties. Summary of the cryo-EM volumes determined in this work, with their properties, for the indicated datasets. Stalk represents the identified stalk conformations^48^. S – straight linker conformation, yellow background; I – intermediate linker conformation, green background; and B – bent linker conformation, blue background.

**Extended Data Fig. 7.**
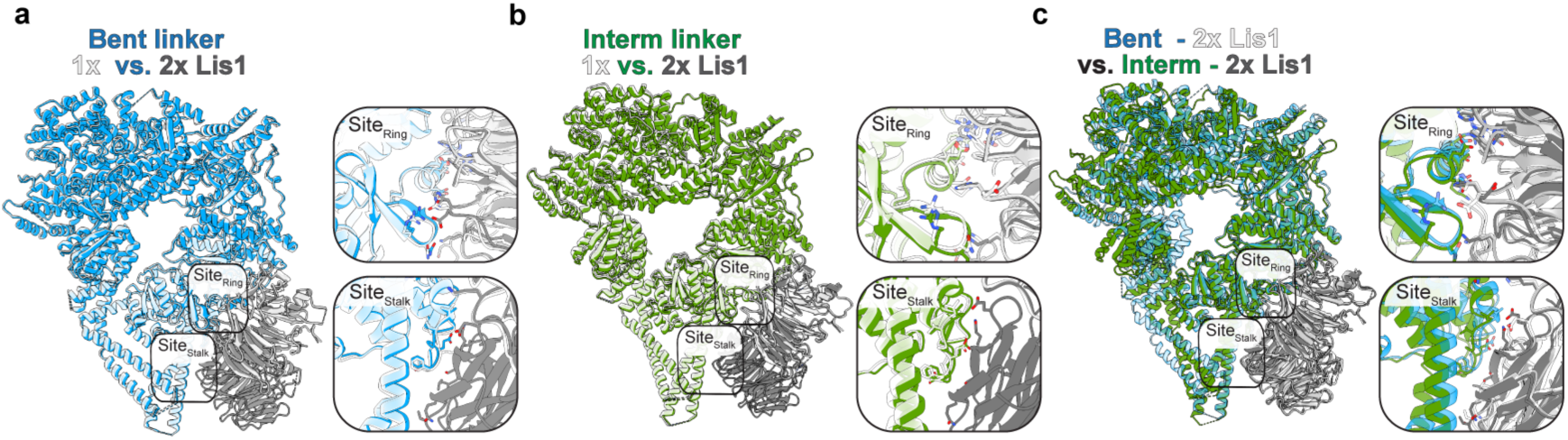
Specifics of Lis1 interaction with dynein. **a**. Comparison of models build for bent linker dynein bound to 2 Lis1 β-propellers (B8, blue, Lis1 – gray) with bent linker dynein bound to 1 Lis1 β- propellers (B7, white, Lis1 – white). The two Lis1 binding sites are highlighted and shown to the right. **b**. Comparison of models build for intermediate linker dynein bound to 2 Lis1 β-propellers (I7, green, Lis1 – gray) with intermediate linker dynein bound to 1 Lis1 β-propellers (I6, white, Lis1 – white). The two Lis1 binding sites are highlighted and shown to the right. **c.** Comparison of models build for bent linker dynein bound to 2 Lis1 β-propellers (B8, blue, Lis1 – white) with intermediate linker dynein bound to 2 Lis1 β-propellers (I7, green, Lis1 – white). The two Lis1 binding sites are highlighted and shown to the right.

**Supplementary Table 1.** S. cerevisiae strains used in this study. *DHA* and *SNAP* refer to the HaloTag (Promega) and SNAP-tag (NEB), respectively. TEV indicates a Tev protease cleavage site. *PGAL1* denotes the galactose promoter, which was used for inducing strong expression of Lis1 and dynein motor domain constructs. Amino acid spacers are indicated by gs (glycine-serine).

| Strain | Genotype | Source |
| --- | --- | --- |
| RPY1 | W303a (*MATa*; *his3-11,15*; *ura3-1*; *leu2-3,112*; *ade2-1*; *trp1-1*) | (Eshel et al., 1993) |
| RPY753 | W303a; *pep4Δ::HIS5*; *prb1Δ*; *P<sub>GAL1</sub>-ZZ-Tev-GFP-3HA-GST-DYN1<sub>331kDa</sub>-gs-DHA*; *pac1Δ::URA3*; *ndl1Δ::CgLEU2* | (Huang et al., 2012) |
| RPY816 | W303a; *pep4Δ::HIS5*; *prb1Δ*; *P<sub>GAL1</sub>-8HIS-ZZ-Tev-PAC1*; *dyn1Δ::CgLEU2*; *ndl1Δ::HygroR* | (Huang et al., 2012) |
| RPY1302 | W303a; *pep4Δ::HIS5*; *prb1Δ*; *PAC11-13xMYC-TRP1*; *P<sub>GAL1</sub>-ZZ-Tev-DYN1<sub>331kDa</sub>*; *pac1Δ::HygroR* | (Toropova et al., 2014) |
| RPY1547 | W303a; *pep4Δ::HIS5*; *prb1Δ*; *P<sub>GAL1</sub>-8HIS-ZZ-Tev-PAC1<sub>R275A,R301A,R378A,W419A,K437A</sub>*; *dyn1Δ::CgLEU2*; *ndl1Δ::HygroR* | (Toropova et al., 2014) |
| RPY1882 | W303a; *pep4Δ::HIS5*; *prb1Δ*; *P<sub>GAL1</sub>-ZZ-Tev-GFP-3HA-GST-DYN1<sub>331kDa</sub>-gs-DHA*; *pac1Δ::URA3*; *ndl1Δ::CgLEU2* | This work |

